# A stimulus-computable rational model of visual habituation in infants and adults

**DOI:** 10.1101/2024.08.21.609039

**Authors:** Gal Raz, Anjie Cao, Rebecca Saxe, Michael C. Frank

## Abstract

How do we decide what to look at and when to stop looking? Even very young infants engage in active visual selection, looking less and less as stimuli are repeated (habituation) and regaining interest when novel stimuli are subsequently introduced (dishabituation). The mechanisms underlying these looking time changes remain uncertain, however, due to limits on both the scope of existing formal models and the empirical precision of measurements of infant behavior. To address this, we developed the Rational Action, Noisy Choice for Habituation (RANCH) model, which operates over raw images and makes quantitative predictions of participants’ looking behaviors in a classic visual habituation paradigm. In a series of pre-registered experiments, we exposed infants and adults to stimuli for varying durations and measured looking time to familiar and novel stimuli. We found that these data were well captured by RANCH. Using RANCH’s stimulus-computability, we also tested its out-of-sample predictions about the magnitude of dishabituation in a new experiment in which we manipulated the similarity between the familiar and novel stimulus. By framing looking behaviors as rational decision-making, this work identified how the dynamics of learning and exploration guide our visual attention from infancy through adulthood.

## 1 Introduction

From birth, humans engage in active learning. Even before they gain independent mobility, infants make choices about what to look at and when to stop looking [22, 44]. Developmental psychologists have both studied this process of visual decision-making as a phenomenon itself and also relied on it as a key methodological tool. Two key phenomena have been especially important: habituation and dishabituation. Habituation occurs when infants show reduced interest upon repeated exposure to the same stimulus, whereas dishabituation entails renewed interest following the subsequent introduction of a novel stimulus. Dishabituation, also known as novelty preference, is often leveraged to infer infants’ perceptual and cognitive abilities [2, 3, 17]: in order for an infant to dishabituate, they must distinguish between the original stimulus and the novel one. Despite the robust documentation of habituation and dishabituation in the literature, the mechanisms underlying these changes in gaze duration remain poorly understood. In this paper, we bridge this gap by introducing a rational model of visual habituation and dishabituation. We show that this rational model quantitatively predicts looking times toward different stimuli in visual habituation paradigms.

An important early conceptual model of infant looking time by Hunter and Ames posited that habituation and dishabituation are influenced by the amount of information to be encoded by the infants [26]. Initially, infants are assumed to look longer at stimuli containing learnable, unprocessed information, showing a familiarity preference shortly after initial exposure (Figure 1A). As exposure accumulates, less information remains unprocessed, leading to shorter looking time at the original stimulus; by comparison, relatively longer looking times to a new stimulus are predicted (i.e. novelty preference). While this model has had significant influence, the absence of specificity regarding the encoding process and a way of measuring the amount of information to be encoded in a stimulus have both permitted the model to be used in post-hoc explanations of looking time measurements. Due to this lack of specificity, researchers can allude to this model to argue that infants fell on the part of the preference curve most consistent with the observed data pattern. This interpretive ambiguity has spurred repeated concerns regarding whether looking time measurements should form the basis for key assertions in developmental psychology [8, 23, 40]. To address this criticism, developmental psychology needs formal models that make quantitative predictions, beyond the qualitative foundations of the Hunter and Ames [26] model.

**Figure 1:**
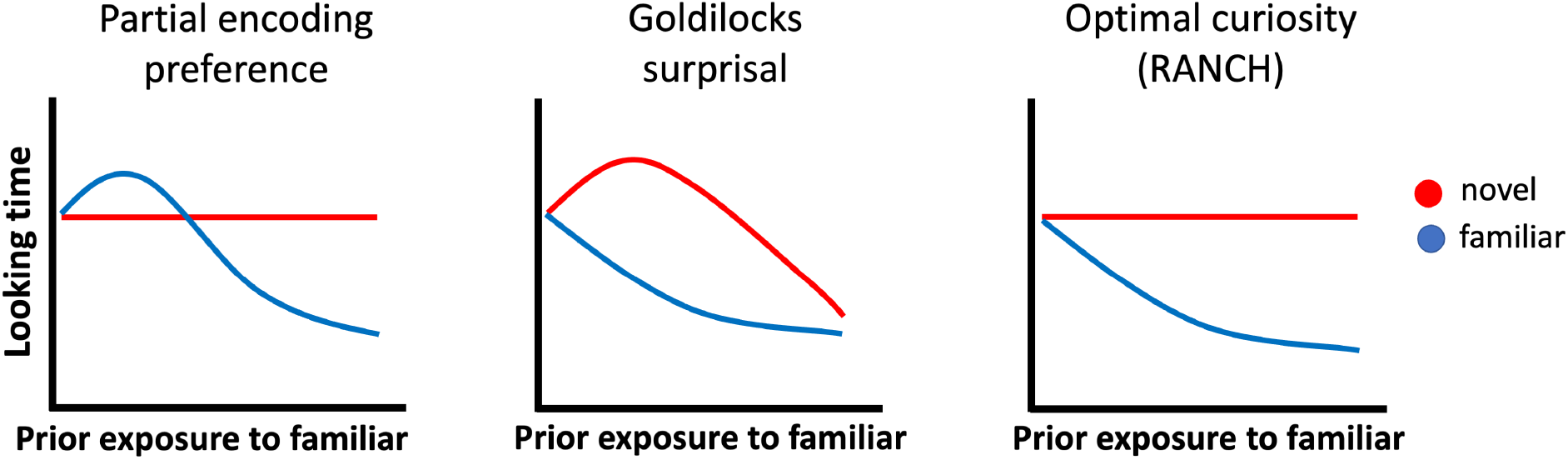
Conceptual models of looking to familiar vs. novel items according to three models of infant attention: (A) The partial encoding model proposed by Hunter and Ames [26] which suggests a shift from familiarity to novelty preferences, (B) The Goldilocks model proposed by Kidd et al [30] which suggests that infants prefer to attend to intermediately surprising events, and (C) the ‘optimal curiosity’ model introduced by Cao et al [10] which suggests that infants are maximizing expected information gain from noisy perceptual samples.

Recent computational models have taken steps towards formalizing the mechanisms underlying looking time changes. One family of models characterizes infants’ looking behaviors through information-theoretic metrics derived from ideal observer models: the models acquire probability distributions from event sequences and derive information-theoretic measures from the ideal learner’s belief before and after each event [30, 41, 42]. For instance, Kidd, Piantadosi, and Aslin [30] found that an ideal learner’s surprisal at an event was related to infants’ looking behavior via a U-shaped curve: infants were least likely to look away when the surprisal of the event was neither too high nor too low (Figure 1B, model predictions derived in Supplementary Information, Figure 8).

While these models provide quantitative fits to variation in looking time, they are not models of how an agent makes the decisions of whether to keep looking at the same stimulus or to look at something else. Rather than producing the target behavior directly, these models describe correlations between model-derived, information-theoretic measures and infants’ overall looking towards whole sequences of events. Relatedly, these models do not explain why infants would ever look at one single stimulus for a long time, rather than encode it instantaneously – there is no account of perceptual noise to be overcome by taking multiple perceptual samples that would justify why looking extends over time [c.f. 9, 29]. Owing to this constraint, these models can only be used to predict infant behaviors in a specific paradigm: looking to a continuous stream of discrete stimuli. Thus, existing models are challenging to generalize to the more common looking time paradigms, in which infants gradually habituate to an individual object or event, and then dishabituate to a new exemplar.

To address these issues, we developed the Rational Action, Noisy Choice for Habituation (RANCH) model [10]. While habituation is a broadly studied phenomenon across cognitive domains – including language acquisition, probabilistic learning, and concept formation – our focus here is on visual habituation, where infants adjust their attention based on repeated exposure to a visual stimulus. Notably, this domain contrasts with the event-based paradigms used in prior modeling work [30, 41, 42], but corresponds well with a substantial portion of the infant habituation literature.

RANCH describes an agent’s looking behavior as rational exploration based on a sequence of noisy perceptual samples. The model construes the looking time paradigm as a series of binary decisions: to keep sampling from the current stimulus, or to look at anything else. The model makes sampling decisions based on the Expected Information Gain (EIG) of the upcoming perceptual sample, choosing to keep looking or look away based on which one would in expectation yield the most information. The RANCH model is therefore a rational analysis of looking behavior [34, 37, 1]: looking is analyzed as optimally adapted to extract information from the environment, given specific constraints on processing (e.g., perceptual noise).

Crucially, RANCH is the first stimulus-computable model of habituation, allowing us to derive quantitative predictions from raw visual stimuli. Previous theoretical accounts have described broad principles of habituation, but they do not generate testable, trial-by-trial predictions of looking behavior. As a result, direct comparisons between RANCH and these models remain challenging: existing models do not specify how an agent decides when to continue looking or disengage, nor do they provide a mechanistic link between stimulus properties and looking time. By explicitly modeling these decision processes, RANCH moves beyond post-hoc explanations and offers a computational framework that can be empirically validated and generalized to new contexts.

We previously validated this rational analysis using a large dataset of adults engaging in a self-paced looking paradigm. We asked adults to look at sequences of stimuli consisting of a repeating familiar stimulus (resulting in habituation), and a single, novel stimulus after varying exposure durations (resulting in dishabituation). Effects of prior exposure and novelty on adults’ looking time were well captured by RANCH [10]. Figure 1C shows schematic predictions of RANCH for the experimental setting described above: variable prior exposure to a familiar stimulus, followed by a novel or a familiar stimulus. Rational exploration of a simple perceptual concept results in a monotonic decrease of attention to familiar items as a function of prior exposure.

Here we directly tested hypotheses about the processes underlying infants’ looking time. To do so, we addressed three challenges. A first critical challenge for differentiating formal theories of looking time in infants is that looking time measurements are typically imprecise estimates, due to low test-retest reliability and small sample sizes [45, 36]. In the current work, we developed a novel, within-subject experimental design that allowed us to sensitively measure infants’ looking time in experiments. In our first experiment, we used this paradigm to measure the dynamics of habituation and dishabituation across different amounts of exposure to a stimulus. Beyond validating RANCH’s predictions in infants, this paradigm enabled us to conduct interpretable developmental comparisons of parameter fits between infants and adults – since we can conduct matched experiments in adults – thereby identifying the processes underlying looking time that may change over development.

A second critical challenge for differentiating theories has been their stimulus-dependency. Researchers can rarely make quantitative predictions about how people would respond to different visual stimuli; even previous computational models have relied on using arbitrary representations that were not directly generated from the stimulus materials [e.g. object location, 30, hand-specified binary feature vector, 10]. To address this issue, we incorporated recent progress in convolutional neural networks (CNNs) to allow RANCH to learn from raw images. The representations of many trained CNNs show striking similarities to how primate brains respond to stimuli, and investigating the representations of CNNs has therefore offered insights into how the visual system encodes objects [14, 25, 53]. The activations of these brain-inspired neural networks form embedding spaces, each of which can be seen as a quantitative hypothesis about how humans embed objects in a lower-dimensional perceptual space [46]. In RANCH, we leveraged these principled stimulus representations to enable RANCH to learn from raw images. In other words, RANCH can ‘see’ an experiment in a way similar to infants, and make predictions for each unique trial.

A final challenge is the generalizability of models to new contexts. Because RANCH can be applied across stimulus sets without additional assumptions, we were able to test its predictions in a new dataset. To conduct these tests, we fit RANCH’s parameters to the dataset from the initial habituation-dishabituation experiment. Then, we used the best-fitting parameters to generate predictions for a new experiment designed to measure differences in dishabituation magnitude. In this experiment, we systematically varied the similarity between the familiar and novel stimulus such that the novel stimulus differed in its pose angle, number, identity, or animacy. This experiment tested a prediction derived from many theories of habituation and dishabituation [26, 19]: that observers’ dishabituation magnitude should be related to the similarity between the habituated stimulus and the novel stimulus. The more dissimilar two stimuli are, the more one should dishabituate to the novel stimulus.

Addressing these three challenges allowed us to empirically test competing hypotheses about habituation and dishabituation using our experimental data (Figure 1). However, because existing models do not generate quantitative predictions, we could not directly compare RANCH to alternative computational models. Instead, we evaluated whether RANCH accurately captured key behavioral patterns in looking time.To preview our results, we found that RANCH provided the best description of behavior in Experiment 1, in which we systematically varied prior exposure to a stimulus before measuring looking time to either the same, familiar stimulus or a novel stimulus. Fitting RANCH to infant and adult data separately and comparing the best-fitting parameters revealed that infants’ learning was tuned for noisier processing than adults. Then, using these best fitting parameters, we generated predictions for a novel task used in Experiment 2 in which we varied the similarity between the familiar and novel stimulus. RANCH generalized to a novel task context by predicting the dishabituation magnitude as a function of stimulus dissimilarity. Together, these model fits provide evidence that habituation and dishabituation to sequential visual stimuli are well described by a rational analysis of looking time. All materials, including the code and data, are available at the project OSF page: https://osf.io/d7jz8/

## 2 Rational Action, Nosy Choice for Habituation model

The Rational Action, Noisy Choice for Habituation model (RANCH) is a Bayesian perception and action model in which a learning model is used to derive optimal perceptual sampling decisions [10]. In RANCH, the learning problem in habituation is conceptualized as learning a visual concept – a single representation underlying a group of observed stimuli. The model learns by observing a series of noisy perceptual samples from each stimulus. During learning, the model makes decisions about how many samples to receive from each stimulus before moving to the next stimulus. We treat the number of samples for each stimulus as being linearly related to looking time duration.

RANCH provides a probabilistic framework for making perceptual decisions. In this section, we will describe the three modular components of RANCH: the perceptual representation, the learning model, and the decision model.

### 2.1 Perceptual representations

We used the deep convolutional neural network (CNN) ResNet-50 to encode the stimuli used in behavioral experiments. ResNet-50 is a standard architecture that is widely used in computer vision as well as in comparisons to human behavior [24, 32]. A total of 50 layers are implemented in this architecture: it starts with a convolution layer, includes several groups of residual blocks that help preserve information through the network, and ends with a global average pooling layer and a fully connected layer for classification. We used a version of ResNet-50 that was pretrained on ImageNet [13]. Training on ImageNet optimizes the ResNet-50’s activations for image classification, and we can use these activations as a perceptual embedding space. When we pass new stimuli through a pre-trained network, we extract these activations to obtain principled embeddings of stimuli (Figure 2A). We use the first three principal components of the embedding space to limit the dimensionality of the downstream learning problem.^1^

**Figure 2:**
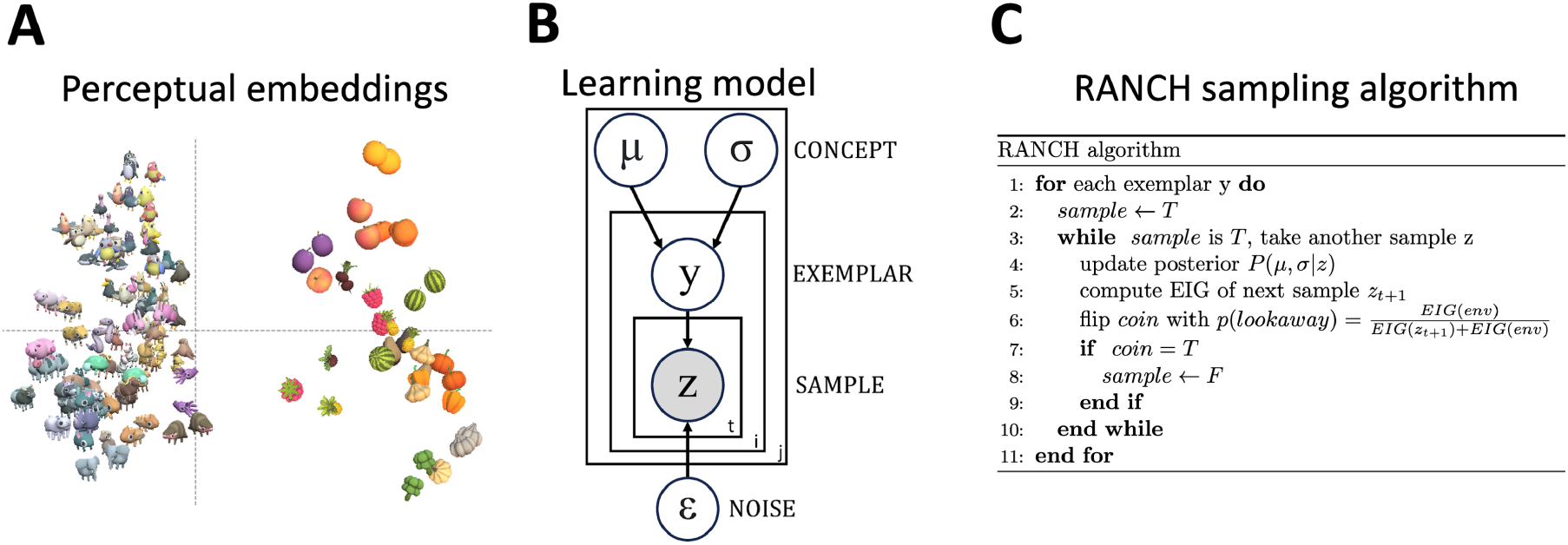
Three components of RANCH: (A) Perceptual representation: we plotted the first two principal components of stimulus embeddings from ResNet-50 final layer activations, (B) Learning model: plate diagram for Gaussian concept learning from noisy observations, (C) Decision model: RANCH repeatedly samples until environmental EIG outweighs EIG from another sample from the same stimulus.

### 2.2 Learning model

RANCH models looking behaviors in habituation experiments as hierarchical Bayesian concept learning [20, 51]. The concept to be learned is a multivariate normal distribution parameterized by *µ, σ*, which represents beliefs about the location and variance of the presented concept in the embedding space. The concept *µ, σ* generates exemplars *y*, which is equivalent to the stimulus seen by the human participants. But RANCH does not directly learn from the stimulus. Instead, it learns from the series of noisy perceptual samples 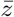 generated by the exemplar. The noisy perceptual sample is corrupted by zero-mean gaussian noise, represented by ϵ. A plate diagram is shown in Figure 2B. At each time step, the model uses one perceptual sample to update its representation of the concept, following Bayes Rule:

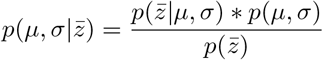

### 2.3 Decision model and linking hypothesis

The Bayesian learning model provides estimates of the target concept based on the observed data, but it does not contain a decision rule for whether to keep sampling from the same stimulus or to move on to the next stimulus. To make this decision, RANCH uses the expected information gain (EIG) of the next perceptual sample. EIG is a metric used under the rational analysis of information-seeking behavior [1, 37], and is computed as the expected value of the KL divergence between the hypothetical posterior at the next timepoint *t* + 1 and the current posterior. Formally: *EIG* = *E*_*t*+1_(*KL*(*P* (*µ, σ* | *z*_*t*+1_), *P* (*µ, σ* | *z*_*t*_))). RANCH then uses a softmax choice (with temperature = 1) between the EIG of the next sample and a constant, which is assumed to represent the amount of information gain provided by the environment when the learner looks away from the stimulus. An algorithm table describing the overall sampling procedure is shown in Figure 2C.

### 2.4 Implementation

RANCH has four free parameters: the learner’s priors over the concept to be learned (*µ, σ*), the learner’s priors over their perceptual noise (*σ*_*ϵ*_) and the constant representing EIG from the environment (*EIG*_*env*_). We conducted a grid search over these parameters.

While simple Gaussian concept learning models can be computed analytically, hierarchical models must be approximated numerically; the addition of perceptual noise to our model puts it in the latter category. We used a grid approximation for inference, creating discrete grids over *µ, σ, y* (exemplars) and *z* (noisy samples) to perform inference. Further, because the EIG computation requires computing an expectation over possible next observations, we approximated this expectation via a grid of possible next observations.

In the simulations below, we ran RANCH for each stimulus ordering. To remove approximation noise resulting from the grid approximations and the stochastic nature of the sampling algorithm, we averaged model results for each stimulus and parameter setting over 100 runs. To avoid bias, these 100 simulations used 20 different random perturbations to the approximation grids such that the exact location of the points in the grids was randomly shifted by small increments. We used the average number of samples the model decided to take from each stimulus as a proxy for looking time.

This process required significant computational resources. The grid approximation, combined with repeated simulations, demanded high-performance cluster GPUs to perform vectorized computation using a PyTorch implementation of our model. Running a single parameter setting through an experiment took approximately 10 hours on a GPU. By distributing different parameter settings across ∼15 GPUs at a time, a full parameter search took about 3 days to complete.

## 3 RANCH predicts habituation and dishabituation in relation to exposure duration

The competing models of habituation and dishabituation shown in Figure 1 make contrasting predictions with regard to the effect of increasing prior exposure on looking time. To directly test these predictions, we collected looking time data from both infants and adults in an experiment where we varied cumulative exposure to a familiar stimulus, and measured subsequent looking time to a novel or familiar stimulus. We then let RANCH “see” the same stimulus sequences used in our behavioral experiments and compared our model’s behavior to that of humans.

### 3.1 Behavioral experiments

#### 3.1.1 Infant experiment

We tested infants in a novel experimental paradigm in which exposure duration to the familiar stimulus was manipulated in a within-subject design shown in Figure 3 (Experiment 1, Infants). Infants were recruited remotely and tested on Zoom video chat software; this testing method increased the efficiency of recruitment and has been shown to yield comparable effect sizes to in-person testing [12, 11]. Each infant saw six blocks. In each block there was an exposure phase and a test trial. During the exposure phase, an animated animal (designated the familiar stimulus) was presented 1 to 9 times. Next, infants’ looking time was measured on a test trial. The test trial showed either the familiar stimulus, or a novel animal. We ran three versions of this experiment, each with three different numbers of exposure events. We tested a combined sample of 103 7-10 month old infants, with 31 infants in the first sample (0, 4 or 8 exposure events), 35 infants in the second sample (1, 3 or 9 exposure events) and 37 infants in the third sample (2, 4 or 6 exposure events; total n=103, *M*_*age*_ = 9.38 months, 47 female).

**Figure 3:**
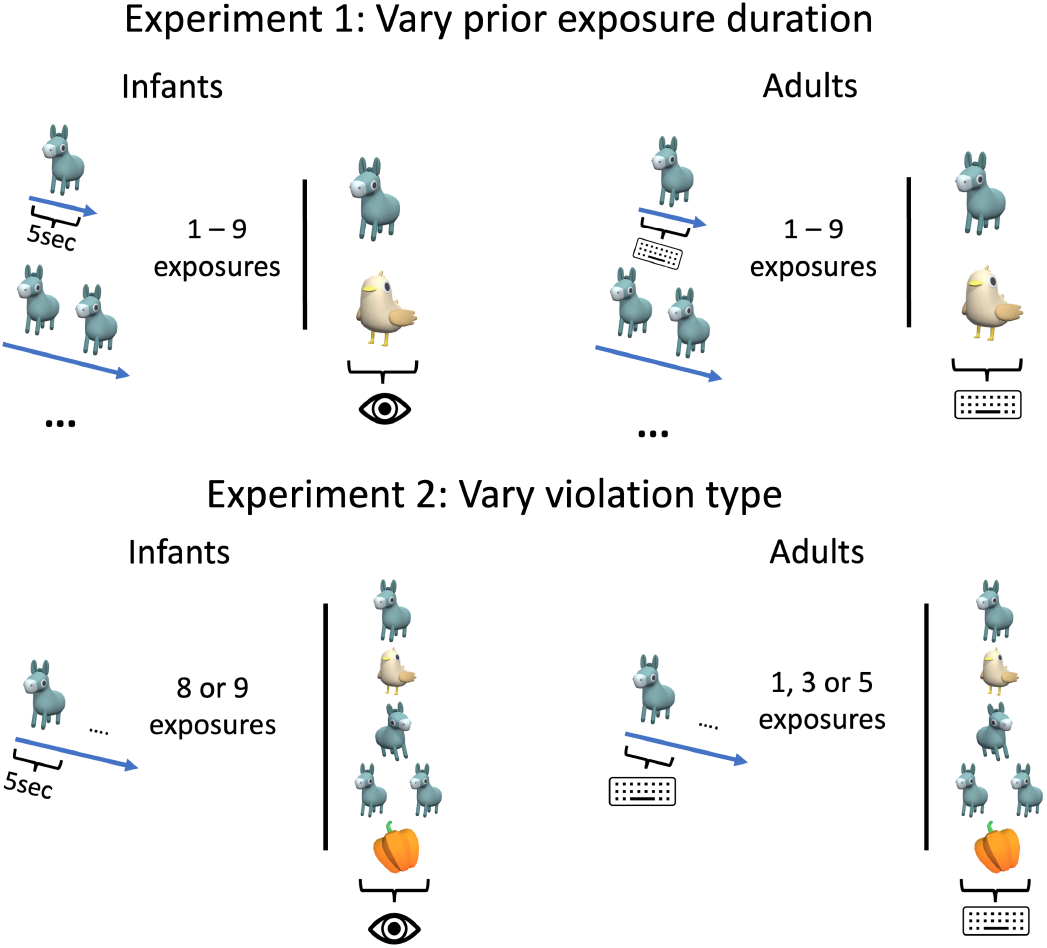
Experimental design for Experiments 1 and 2, infants and adults. The vertical line separates the exposure phase and the test trial. Experiment 1 varied exposure to a stimulus and measured looking to familiar or novel stimuli. Experiment 2 varied how the familiar stimulus was violated, by changing the pose, identity, number or animacy of the stimulus. Infants had fixed prior exposure durations (5 second per exposure) and looking was measured until a 2-second lookaway. Adults responded on each trial via a keypress to continue to the next trial.

#### 3.1.2 Adult experiment

We ran a self-paced looking time experiment on Prolific, collecting data from 470 adult participants (Experiment 1, Adults; see Figure 3 for design). Stimuli used in the adult experiment were drawn from the same set used in the infant experiment. During the experiment, participants were asked to watch sequences of stimuli at their own pace, and they could press a key to proceed to the next stimulus whenever they wanted to. The time between the onset of the stimulus and the key press was used as a proxy for looking time. Each sequence began with a familiar stimulus repeating 1 to 10 times, followed by either the same familiar or a novel stimulus.

#### 3.1.3 Modeling Procedure

RANCH received the ResNet-50-derived stimulus embeddings as input, presented in sequences mirroring the structure in our infant and adult experiments (Figure 3). To simulate a block of the infant experiment, the model was first presented with the “familiar” stimulus. For each stimulus, the model took 5 samples each and then moved on to the next stimulus. This phase was designed to be the exposure phase infants went through, since during the exposure phase infants had no control over the progression of the stimulus sequence. After the exposure phase, the model was presented with the test trial, which was either familiar or novel. We used the number of samples the model took on this test trial as a proxy for infant looking time. To simulate the self-paced adult experiment, we computed the number of samples taken by RANCH on every trial in each stimulus sequence.

### 3.2 Behavioral and model results

The paradigm successfully captured habituation in both infants and adults. In infants, as the number of trials in exposure phase increased, the looking time at the familiar stimulus in the test trials decreased (*β* = -4.55; *SE* = 0.92; *t* = -4.92; *p* < 0.001). Infants clearly dishabituated on trials with longer exposures, looking longer at the novel stimulus than the familiar stimulus after long exposure (8 exposures: *β* = 0.82; *SE* = 0.22; *t* = 3.81; *p* < 0.001, 9 exposures: *β* = 0.51; *SE* = 0.12; *t* = 4.21; *p* < 0.001). The adult experimental paradigm also successfully captured habituation and dishabituation (Trial number: *β* = -0.01; Trial Type: *β* = 0.22; Both *SE* < 0.001; Both *p* < 0.001). We found no evidence for familiarity preferences or non-linearities in the shape of the dishabituation curve in infants or adults (cf. Figure 1A&B).

RANCH predictions qualitatively matched habituation and dishabituation in both infants and adults. To quantitatively evaluate these predictions, we fit a linear model (adjusting model-generated samples by an intercept and scaling factor) and then assessed two complementary metrics. First, the root mean squared error (RMSE) captures the absolute error in the same units as looking time. Second, the coefficient of determination (*R*^2^) measures the relative variation in looking time that is explained by the scaled model predictions. Since each metric relies on different assumptions and highlights distinct aspects of predictive accuracy, they together provide a more robust assessment of model performance. We minimized overfitting by employing cross-validation – using a split-half design for infant data and ten-fold for adult data – to compute both RMSE and *R*^2^ on held-out samples. Cross-validation provides numerical estimates of out-of-sample generalization performance that can be used for numerical model comparison.

**Table 1:** Cross-validated RMSE across linking hypotheses and lesioned models. Mean RMSE and *R*^2^ is averaged across all parameter values, best RMSE and *R*^2^ is the single best-performing parameter set.

| Linking Hypothesis | Infants |  |  |  | Adults |  |  |  |
| --- | --- | --- | --- | --- | --- | --- | --- | --- |
| | Mean RMSE | Mean $R^2$ | Best RMSE | Best $R^2$ | Mean RMSE | Mean $R^2$ | Best RMSE | Best $R^2$ |
| EIG | 3.939 | 0.330 | 3.696 | 0.457 | 0.222 | 0.883 | 0.166 | 0.905 |
| KL | 3.912 | 0.339 | 3.658 | 0.386 | 0.224 | 0.882 | 0.163 | 0.899 |
| Surprisal | 4.396 | 0.179 | 3.601 | 0.465 | 0.303 | 0.647 | 0.213 | 0.845 |
| No learning | 4.463 | 0.067 | 4.135 | 0.295 | 0.449 | 0.037 | 0.377 | 0.304 |
| No noise | 4.224 | 0.198 | 3.875 | 0.323 | 0.339 | 0.504 | 0.292 | 0.615 |

### 3.3 Comparison with alternative linking hypotheses

Here we evaluated different ways of specifying RANCH’s decision-making mechanism (i.e., different “linking hypotheses” within RANCH). EIG is the optimal method for making sampling decisions [35]. We compared this method to two other information-theoretic measures proposed in prior literature: Kullback-Leibler divergence (KL-divergence) and surprisal. KL-divergence measures the information gained when one’s beliefs are updated, effectively quantifying “learning progress”, while surprisal captures how inconsistent incoming information is with prior expectations. Both have been shown to be associated with infants’ looking behaviors [30, 41]. We then repeated the cross-validation process using these linking hypotheses and compared the model’s performance based on the parameter that provides the best average fits to the test sets according to RMSE. Out of the three linking hypotheses, we found that EIG and KL-divergence were comparable to each other but surprisal-based RANCH performed worse in both populations (Table **??**). For both infants and adults, this poorer fit seems to have been primarily driven by the mismatch between our participants’ and RANCH’s looking to familiar stimuli (blue curves, Figures 4 and 5).

**Figure 4:**
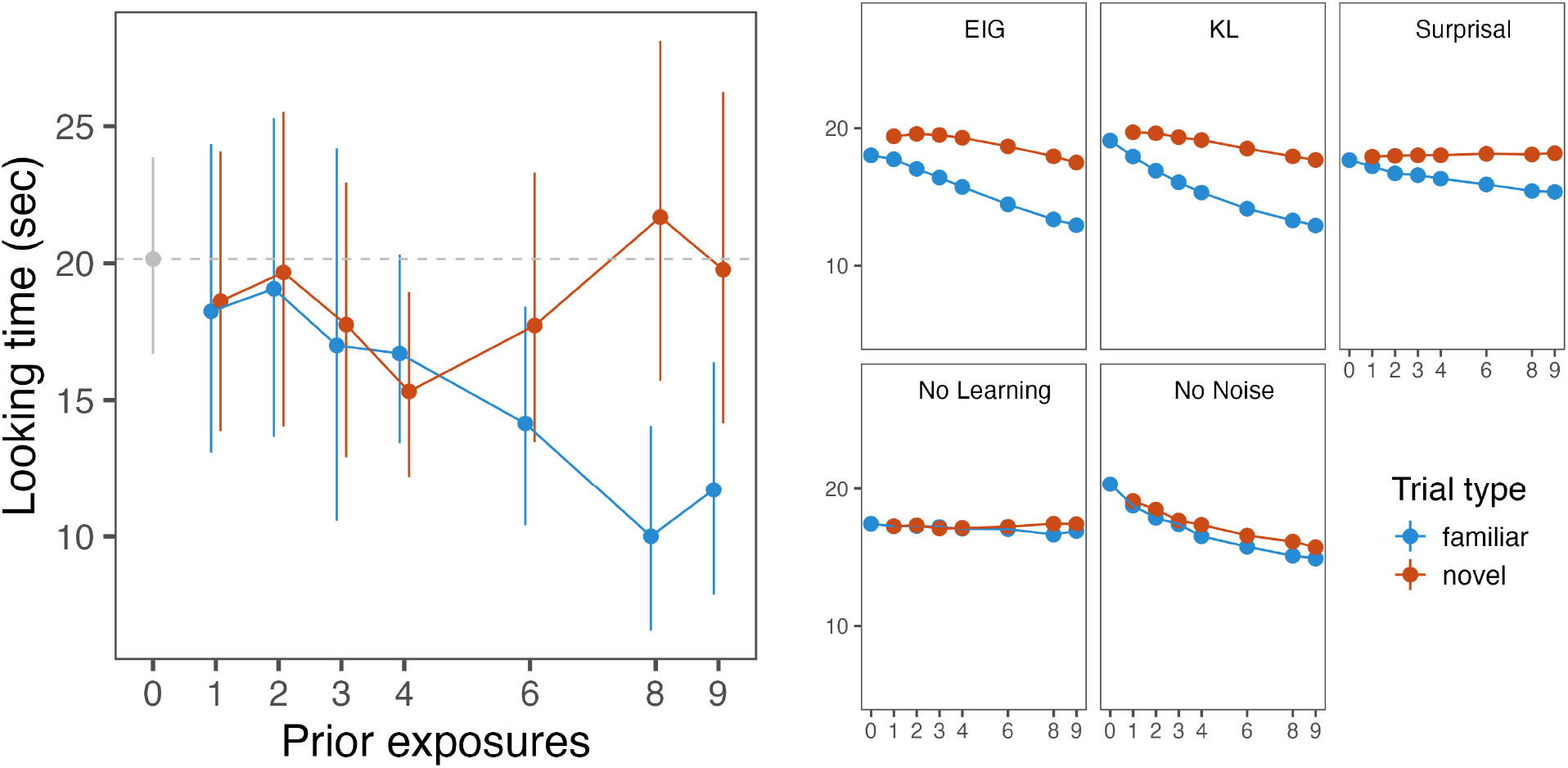
Experiment 1, Infant and RANCH behavior (different linking hypotheses and lesioned models). The x-axis shows the number of prior exposures shown to infants and RANCH before measuring looking time / number of samples on the test trial, which was novel or familiar. The y-axis shows the mean looking time in seconds for the behavioral panel, and the scaled model samples for RANCH. The scaling procedure was applied to each linking hypotheses/lesioned model separately. We found evidence for habituation and dishabituation (after long prior exposures), but no evidence for familiarity preferences. Error bars show the standard error of the mean.

**Figure 5:**
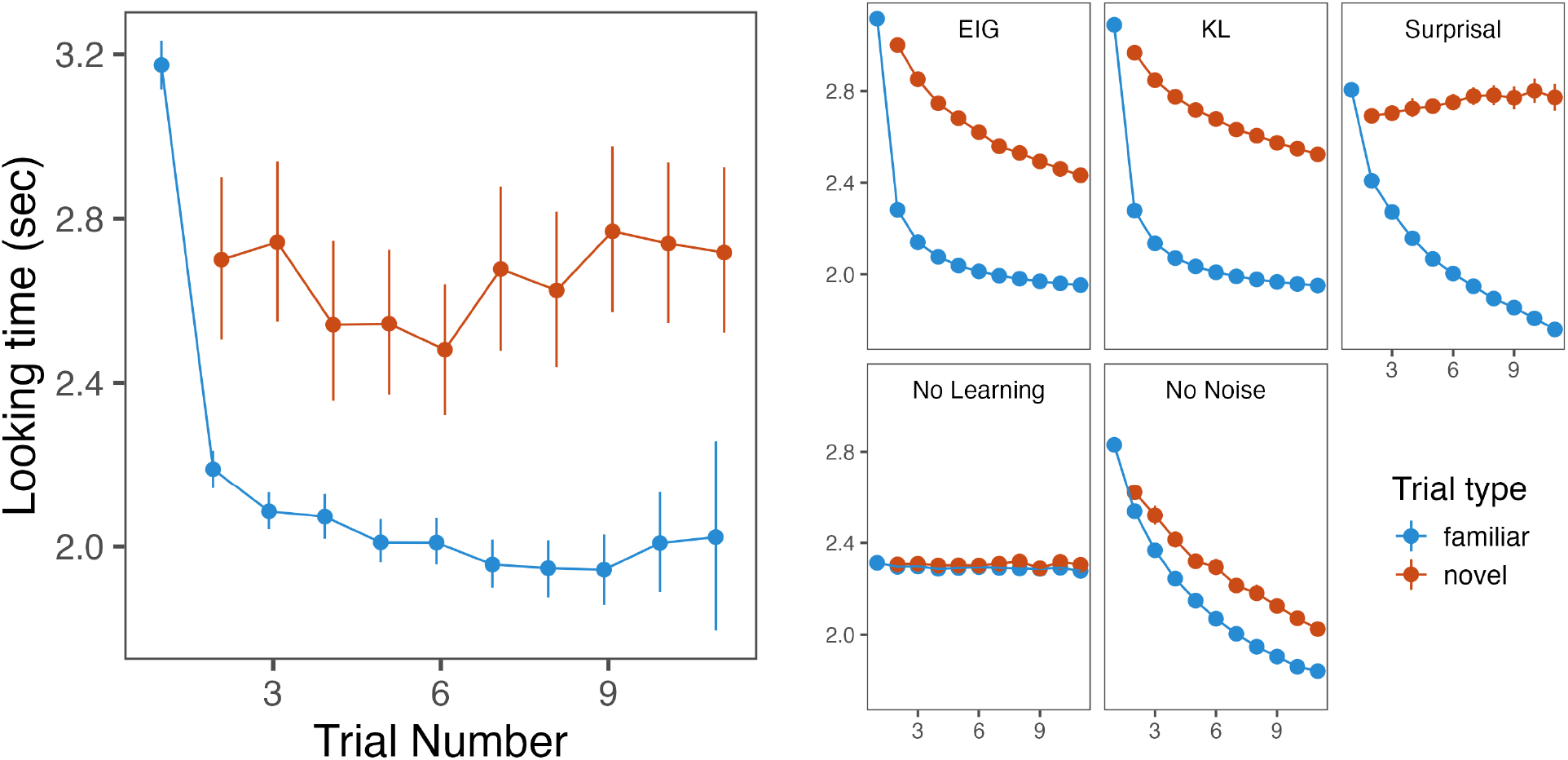
Experiment 1, Adult and RANCH behavior (different linking hypotheses and lesioned models). For adults and the corresponding RANCH simulations, looking time was measured on every trial, shown on the x-axis. The y-axis shows the mean looking time in seconds for the behavioral panel, and the scaled model samples for RANCH. Similar to infants, we found evidence for habituation and dishabituation, but no evidence for familiarity preferences. Error bars show the standard error of the mean.

### 3.4 Comparison with lesioned model

We tested the importance of two of RANCH’s key components by creating lesioned models. In the first lesioned model, we removed noisy sampling by setting the learner’s noise prior, *σ*_*ϵ*_, to 0 (“No noise”). In the second lesioned model, we removed the learning model, and replaced the EIG-based sampling with a random sampling policy (“No learning”). Both lesioned models provided poor fits to the behavioral datasets (Table **??**). This suggests that both noisy perception and the learning model are critical to RANCH’s performance.

### 3.5 Parameter interpretation and developmental comparison

One advantage of RANCH is the interpretability of its parameters. Beyond finding that RANCH’s fit to the data is robust across parameters, we can also ask which specific parameters maximize that fit. In particular, comparing the best fitting parameters between infants and adults could provide insights into the origin of the developmental differences in behavior (see “Joint Scaling” under Methods for technical details). When investigating which parameters were most influential in achieving better fit, we found that *σ*_*ϵ*_, the perceptual noise parameter, played a key role in improving fit. A high *σ*_*ϵ*_ improved fit to the infant data, but a low *σ*_*ϵ*_ improved fit to the adult data (Supplementary Information, Figure 11). Conceptually, *σ*_*ϵ*_ represents an agent’s beliefs about how much noise there is in their own perceptual input; the difference in *σ*_*ϵ*_ suggests that infants may have adjusted their learning behavior to higher perceptual noise.

### 3.6 Discussion

Using within-subject measurements of looking time, we examined the sensitivity of looking to familiar vs. novel stimuli as a function of prior exposure. We did not find evidence for non-linearities predicted by prior models (Fig. 1A & 1B), and instead found that habituation and dishabituation followed monotonic trends as predicted by RANCH (Fig. 1C). Although these non-linearities could in principle still remain, our experiments had substantially larger samples and more graded manipulations of exposure duration than previous experiments with infants or adults. Next, we tested whether RANCH can make predictions in a novel looking time experiment.

## 4 RANCH predicts stimulus similarity-based dishabituation in infants and adults

Beyond predicting non-linearities in attentional preferences during encoding (Fig. 1B), Hunter and Ames [26] also predicted that after complete encoding of a stimulus, dishabituation magnitude should be linked to the similarity between two stimuli: the more dissimilar the novel stimulus is to the habituation stimulus, the more participants should dishabituate. This prediction is important theoretically because it underlies the use of looking time as a continuous measure of stimulus similarity (as in, e.g., [49]).

We designed an experiment testing this prediction by systematically varying the similarity between the familiar and novel stimuli within each block. RANCH provides an explicit notion of similarity through the use of ResNet50-derived image embeddings. We can therefore test whether RANCH also predicts such similarity-based differences in dishabituation magnitude. We further tested RANCH by assessing whether the best-fitting parameters in our previous experiment can be used to predict infants’ and adults’ looking time in a new experiment.

### 4.1 Behavioral Experiment

#### 4.1.1 Infant experiment

To study whether infants’ dishabituation was modulated by similarity between the habituation and novel stimulus, we ran an experiment, again recruiting via Zoom (n = 57, age range = 7-10 months, *M*_*age*_ = 8.99 months). We also ran an exact replication via Children Helping Science (CHS), an asynchronous online platform for online developmental data collection (Scott and Schulz [47]; n = 66, age range = 6-12 months, *M*_*age*_ = 9.69 months). We presented infants with a series of 4 blocks, each again consisting of a exposure phase and a test trial (Figure 3, Experiment 2, Infants). Instead of modifying prior exposure as in the previous experiment, here infants were always familiarized 8 or 9 times with an animal. We chose 8 and 9 exposures based on the results of Experiment 1, where these durations caused robust dishabituation. Moreover, using slightly variable exposure durations reduces the risk that infants develop fixed expectations about when a novel stimulus will appear. After the exposure phase, infants saw a test trial, which could relate to the familiar animal in one of five ways: they either saw the familiar animal again (familiar), the same animal facing in the opposite direction (pose), a different animal facing the same way (identity), two instances of the same animal side-by-side (number), or an inanimate vegetable or fruit (animacy).

In a separate experiment, we also tested baseline interest in these stimuli without prior exposure, to ensure that condition differences in the main experiments were due to violations, rather than intrinsic to the stimuli (n = 35, age range = 7-10 months, *M*_*age*_ = 9.21 months). This experiment revealed no baseline differences in looking time between categories (Supplementary Information, Figure 13).

#### 4.1.2 Adult experiment

We tested adults in an experiment that was procedurally similar to Experiment 1 but whose stimuli were similar to the current infant experiment. Participants (N = 468) saw 24 blocks of animations, each of which was either 2, 4 or 6 trials long, and pressed a button to continue to the next trial in the sequence. 8 blocks showed a single, familiar stimulus repeatedly. 16 blocks always began with one stimulus shown 1, 3 or 5 times, and ended with another, novel stimulus. Unlike the infant experiment, which always started with a single animal, the adult experiment was long enough to show all permutations of familiar vs. novel blocks: Sequences could start with any stimulus type, i.e. either animals or vegetables, facing either left or right, presented either singly or in a pair. The novel stimulus then would violate one of the properties (pose, identity, number or animacy).

### 4.2 Behavioral and model results

#### 4.2.1 Infant results

In the infant experiments, we found similar results for the samples tested synchronously over Zoom and asynchronously via CHS. For the Zoom sample, We fit a mixed effects model to predict log-transformed looking times from the violation type, and found that infants dishabituated to all novel test trials (identity: *β* = 0.33; *SE* = 0.16; *t* = 2.08; *p* = 0.04, number: *β* = 0.32; *SE* = 0.17; *t* = 1.92; *p* = 0.057, animacy: *β* = 0.33; *SE* = 0.16; *t* = 2.02; *p* = 0.046), except the pose violation, which elicited no significant dishabituation (pose: *β* = 0.13; *SE* = 0.16; *t* = 0.83; *p* = 0.406).

We applied the same mixed effects model to the CHS sample. We again found that animacy and number resulted in the strongest dishabituation in infants (animacy: *β* = 0.34; *SE* = 0.14; *t* = 2.47; *p* = 0.015, number: *β* = 0.35; *SE* = 0.13; *t* = 2.68; *p* = 0.008), and pose violations did not elicit dishabituation (*β* = 0.02; *SE* = 0.13; *t* = 0.14; *p* = 0.888). However, unlike the Zoom study, the identity violation did not elicit significant dishabituation in our CHS study (*β* = 0.07; *SE* = 0.14; *t* = 0.46; *p* = 0.646).^2^

We then combined the two infant datasets (total n = 123) to gain additional power. In an exploratory analysis, we ran the same linear model as on the individual datasets, since we found no additional variance explained by a random effect of dataset. We again found significant dishabituation to animacy and number violations (number: *β* = 0.34; *SE* = 0.11; *t* = 3.19; *p* = 0.002, number: *β* = 0.35; *SE* = 0.1; *t* = 3.37; *p* = 0.001), marginally significant dishabituation to identity violations (*β* = 0.19; *SE* = 0.11; *t* = 1.77; *p* = 0.078) and no dishabituation to the pose violation (*β* = 0.07; *SE* = 0.1; *t* = 0.65; *p* = 0.514). Together, we found evidence that the magnitude of dishabituation was related to the similarity of the novel trial, as infants’ looking time followed our expected ordering: animacy > number > identity > pose > familiar (Figure 6).

**Figure 6:**
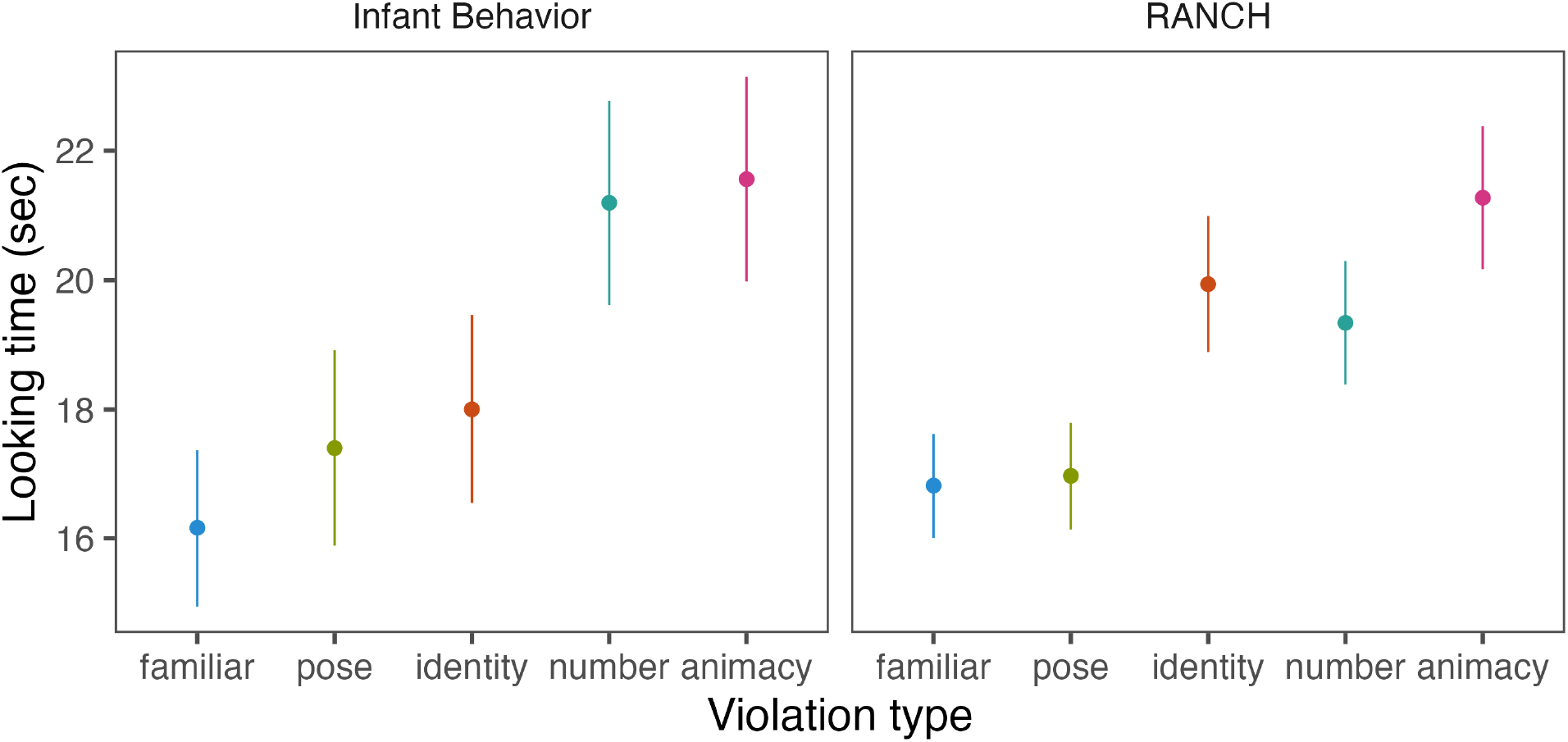
Experiment 2, Infant and RANCH behavior. The x-axis shows how the familiar animal was violated and the y-axis shows the mean looking time in seconds / scaled model samples for RANCH during test trials following the exposure phase. Error bars show the standard error of the mean. Both infants and RANCH showed a graded pattern of dishabituation depending on the violation type. We show the results using the combined dataset of the Zoom and Children Helping Science experiments.

Next, we tested whether infants’ dishabituation was better described by a “same or different” account, in which dishabituation responds similarly to any novel stimulus, or a “graded” account, in which the type of violation influences the magnitude of dishabituation. To do so, we conducted a model comparison in which violation type was either parsed into two levels (familiar or novel) or five levels (familiar, animacy, number, identity or pose). The model comparison significantly favored the more granular model (*χ*(3) = 8.9; *p* = 0.031), suggesting that infants were sensitive to the type of violation rather than just to whether the test stimulus was familiar or novel.

#### 4.2.2 Adult results

In adults, as in previous experiments, we found evidence for habituation (Figure 7; *β* = -0.02, *SE* = 0, *p* < .001). More importantly, we also found evidence for graded dishabituation: the dishabituation magnitude to animacy violations was larger than to number violations (*β* = 0.17, *SE* = 0.04, *p* < .001) and to pose violations (*β* = 0.18, *SE* = 0.04, *p* < .001). The same pattern was found for identity violations (*β* = 0.18, *SE* = 0.04, *p* < .001). However, animacy violations were not different from identity violations, nor were number violations different from the pose violations (all p > 0.1). Qualitatively, we found that the ordering for dishabituation magnitude was animacy > identity > pose > number (Figure 7).

**Figure 7:**
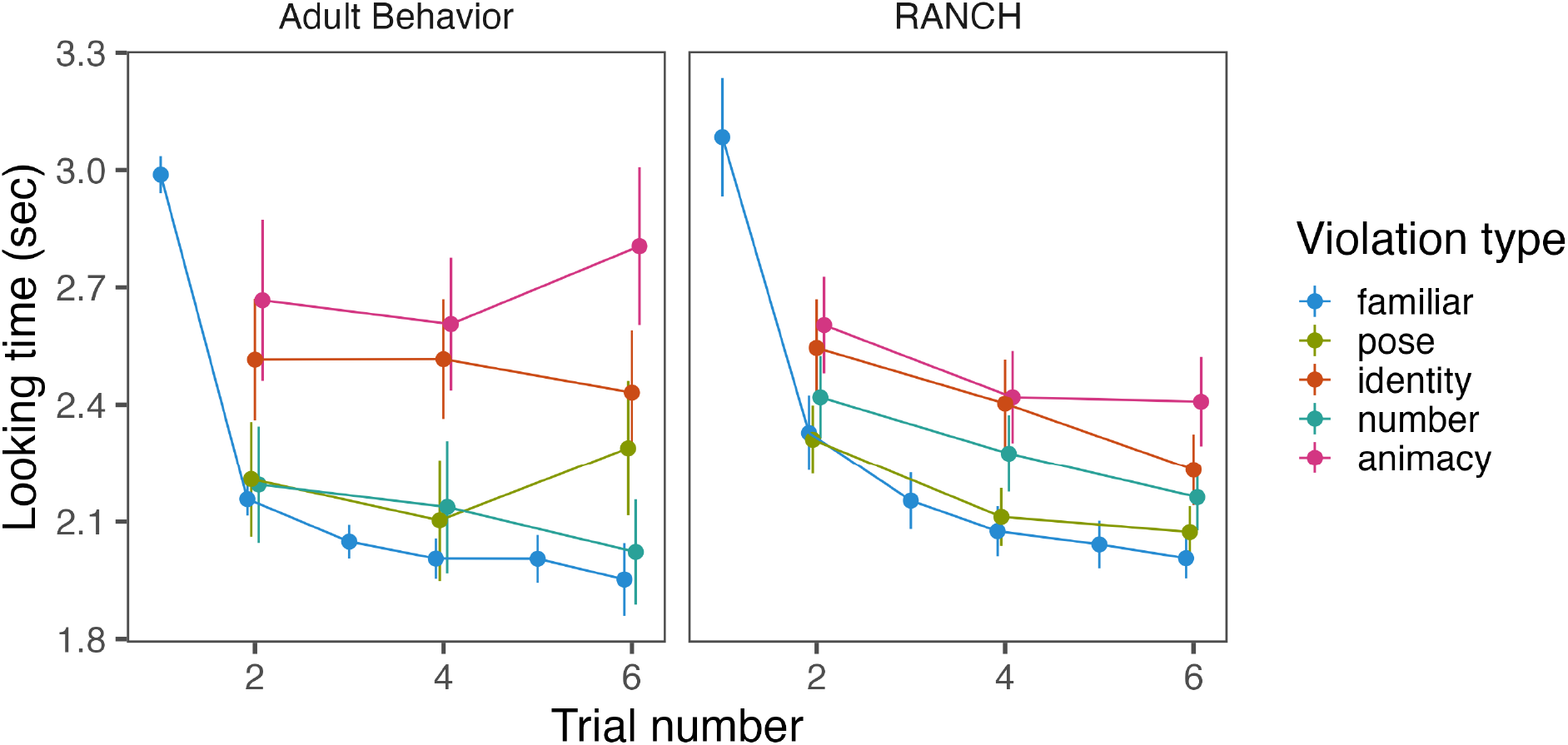
Experiment 2, Adult and RANCH behavior. The x-axis shows the position in the block (which could be of length 2, 4 or 6), and y-axis shows looking time/scaled model samples. Both adults and RANCH showed a similar pattern of graded dishabituation in this task. Error bars show the standard error of the mean.

**Figure 8:**
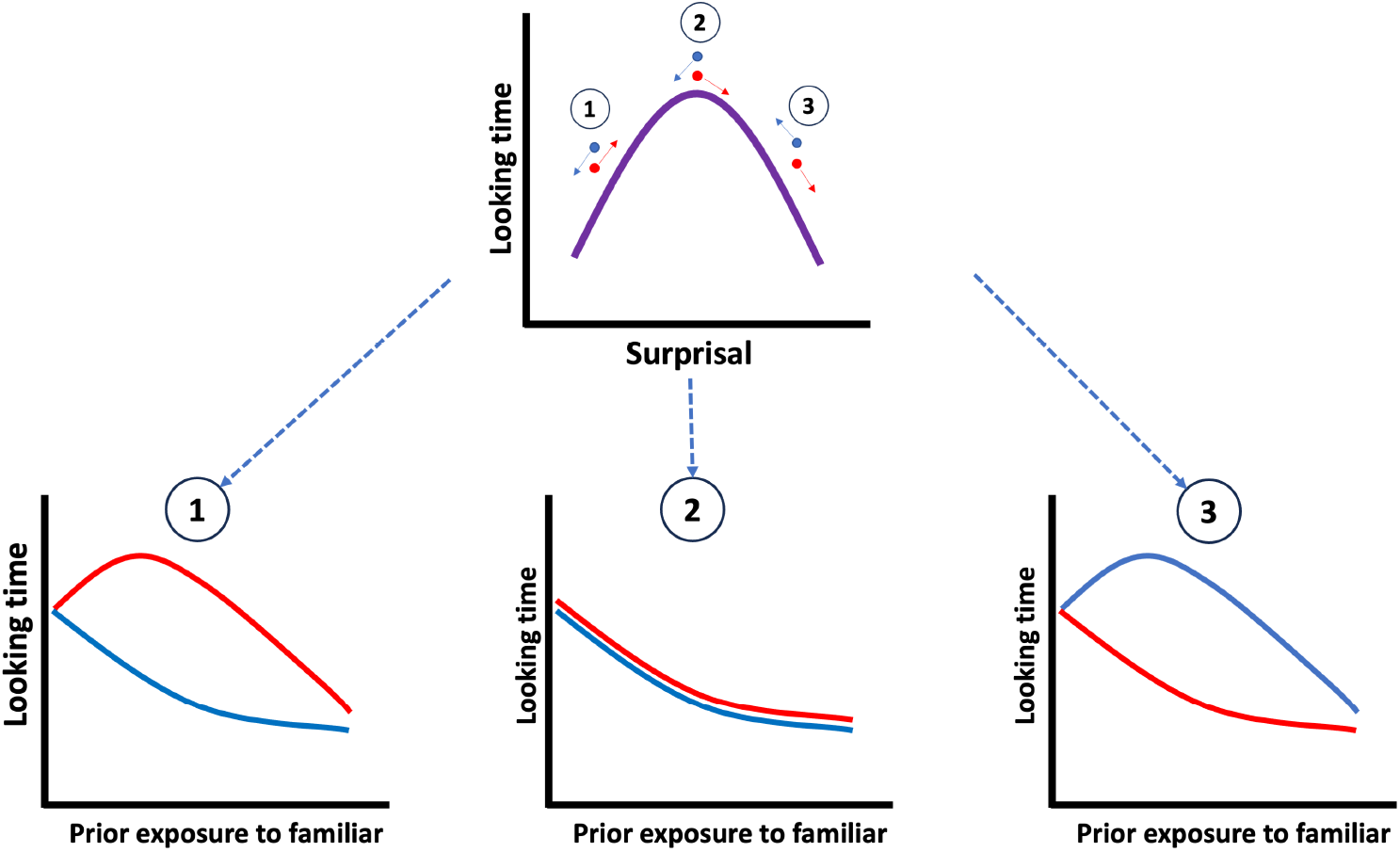
Illustration of how predictions from the Goldilocks model in Fig. 1 were derived.

#### 4.2.3 Model results

We next tested whether RANCH made similar predictions. We used the modeling procedures for the infant and adult experiments described above.

Using the best-fitting parameters from Experiment 1, we found that RANCH generalized to Experiment 2 successfully (Infants: *RMSE* = 1.26; *R*^2^ = 0.66; Adults: *RMSE* = 0.16; *R*^2^ = 0.72). Moreover, RANCH also showed a qualitative ordering of the dishabituation magnitude that was similar to those seen in infants and adults: animacy > identity > number > pose. This ordering was in line with the ResNet-50-derived embedding distances between the different stimulus categories (Figure 12).

Taken together, these data suggest that RANCH is not only capable of capturing habituation and dishabituation, but also can generalize to a new task and make predictions about dishabituation magnitude in relation to stimulus similarity. We did observe some differences in the relative ordering of dishabituation magnitudes between infants, adults, and RANCH – we discussed these in more depth in the General Discussion.

## 5 General discussion

Even though habituation has been a key method for gaining insight into the infant mind, the underpinnings of this behavior are not well understood. Here, we investigated how well a specific form of visual habituation and dishabituation can be understood as rational exploration of noisy perceptual samples. To do so, we introduced the RANCH model, which takes noisy samples from stimuli and makes decisions about whether or not to continue sampling based on the expected information gain of a new sample. We used RANCH to make predictions about infants’ and adults’ looking time across a range of exposures, and the model showed good fit across a range of parameters. In a second experiment, we found that the same parameters in RANCH could be used to make predictions about looking time in a new task where we varied the similarity between the familiar and novel stimulus. These results suggest that RANCH provides a quantitative account of looking time as resulting from optimal exploration of noisy perceptual samples.

Our work successfully addressed the three critical challenges in testing models of infant looking time: imprecise measurement, stimulus-dependency, and poor generalizability. First, our novel within-subject experiment design, combined with online testing and automatic gaze coding, allowed us to sensitively measure the infant looking time in a relatively large sample. Second, our model leveraged principled stimulus representations to learn from raw images, overcoming the stimulus-dependency of previous models. Lastly, our model can be applied across different stimulus sets, and therefore be used to make out-of-sample predictions about new experiments.

This current work focuses on visual habituation, a fundamental but specific form of habituation that applies to sequential visual stimuli. While habituation has been studied across various domains, our model is specifically designed to account for looking time changes in response to repeated visual exposure. This focus aligns with our choice of perceptual representations derived from CNNs, which process visual inputs rather than abstract probabilistic structures. Visual habituation plays a foundational role in infant cognition, as it provides a mechanism for concept learning based on visual experience. However, it does not encompass all forms of habituation, particularly those involving complex rule learning, linguistic structures. Similarly, RANCH does not capture more global attention dynamics, such as block-to-block attentional drift independent of stimulus properties. Future work should investigate whether models like RANCH can be extended to capture habituation mechanisms in broader learning contexts.

### 5.1 Unique strengths of the RANCH framework

A key strength of RANCH is the cognitive interpretability of its parameters. RANCH integrates structured perceptual representations with Bayesian inference, allowing for stimulus-computable predictions of looking behavior and interpretable parameters at the same time. This integrated approach has been used to study selective attention [7]. Moreover, previous Bayesian models of infant attention have leveraged parameter interpretability to draw conclusions about the computational mechanisms underlying individual differences in attention [43]. Here, we used parameter fits of RANCH to study population differences: By applying a “joint scaling” procedure in which we scaled model samples to infant and adult simultaneously, we could identify which parameters had to be different between infant and adults to account for their different patterns of looking time. Using this procedure, a key developmental difference was in *σ*_*ϵ*_, the learner’s prior on their perceptual noise. Given that *σ*_*ϵ*_ was preferred to be higher in infants than in adults, this suggests that infants’ learning in our experiments may be tuned for noisier perceptual processing. This finding is consistent with previous works showing that infants’ visual contrast sensitivity is indeed constrained by higher level of neural noise [48, 27].

Another strength is RANCH’s ability to model looking on a more fine-grained timescale. Prior research has largely resorted to modeling on the trial level, and asking about correlations between behavior on a given trial (e.g. looking time) and information-theoretic properties of that trial relative to prior training, such as KL divergence or surprisal [30, 42]. Rational analysis relies on computing expected information gain. However, the resolution of these prior models does not support calculating this metric on sample-by-sample basis, since they carve up time only in terms of trials, rather than samples. In contrast, the sample-based modeling framework used in RANCH allows for implementation of the full rational analysis [37, 1], in which learners ask “How much information do I expect to gain from continuing to look at the stimulus?”. The ability to conduct the rational analysis enhances our understanding of habituation as optimally adapted to environmental constraints [see e.g. 19, for a different rational analysis of habituation]. Still, replacing EIG with information-theoretic proxies in our setting can help us examine whether participants’ behavior seems to follow an EIG-based, or a proxy-based, linking hypothesis. In our analyses we found that KL divergence performed comparably to EIG, and the surprisal-based model was worse at fitting both infant and adult data in Experiment 1.

Given that EIG is defined as the expected value of KL-divergence, this suggests that our participants’ strategy is consistent with a “learning progress” metric when making attentional decisions, either backward-looking (KL) or forward looking (EIG) [21]. A surprisal-based strategy however is less consistent with our data, in contrast with other models which relate infant looking to a form of surprise [49, 30].

More generally, while there is no single threshold for what constitutes a “good” model fit, the strength of our approach lies in the relative comparisons across model variants, linking hypotheses, and ablation studies. In this way, we treat model fit not as an absolute benchmark, but as an empirical tool to adjudicate among alternative explanations and assess the mechanistic plausibility of the model’s components.

### 5.2 Within-subject design reveals graded dishabituation

To test our model, we developed novel paradigms for within-infant measurements of habituation and dishabituation. These novel designs allowed us to go beyond testing simple qualitative predictions (i.e. longer versus shorter looking duration). Instead, we validated RANCH’s continuous predictions about the relationship between prior exposure and looking to familiar vs. novel stimuli (Experiment 1) and between perceptual distance and looking (Experiment 2), within subjects. It is rare to observe such graded differences in looking: Most looking time studies measure a simple contrast between familiar and novel, or expected vs. unexpected stimuli and therefore do not capture nuances in dishabituation magnitude as a function of experimental parameters [for an exception see; Téglás et al. [49]]. Our findings revealed quantitative nuances of looking time patterns and provide a higher bar for fitting computational models to such data.

However, Experiment 2 also revealed interesting discrepancies across age groups and RANCH, particularly for number violations: The number of samples RANCH took from number violations was less than identity violations but more than pose violations. In contrast, adults looked at the number violations less than the pose violations, and infants looked at number violations more than identity and pose violations. This discrepancy between the model and adults on the one hand, and infants on the other hand could be explained by different underlying representations for number.

RANCH is designed for representing single objects, not multiple objects, therefore any dishabituation to number violations resulted primarily from simple, low-level perceptual distances, rather than dishabituation to number per-se. To the extent that the dishabituation observed in the data exceeded RANCH’s, this points to other, conceptual sources of new information that cannot be explained purely by low-level visual differences. This is consistent with infant literature showing that infants are sensitive to changes in number specifically, rather than visual changes generally [52, 18]. Adults on the other hand did not show this exaggerated effect of number violations, possibly because adults’ processing was shallower in our task, which is supported by their shorter reaction time.

### 5.3 No evidence for familiarity preferences

A key feature of our experimental data, as well as our model, is that at no point, even for the shortest exposure durations, did we observe longer looking to familiar than to novel stimuli. This finding contrasted with predictions made by a prominent model of infant attention (Figure 1A), which posited that at intermediate levels of encoding, familiar stimuli should be more interesting than novel stimuli. The intuition is that some initial familiarity with a stimulus may allow a viewer to “break into” the information there is to gain from the stimulus – a completely novel stimulus may be too complex or foreign for extended attention to be worth it. However, in the learning setting we used here, neither the rational analysis nor data from human learners showed this strategy. This result suggests at least that limited exposure does not generically lead to familiarity preferences in all experimental paradigms.

The absence of familiarity preferences in our results does not rule out their existence in general. Familiarity preferences may be more subtle than novelty preferences, so that the statistical power that is needed to find familiarity preferences is higher than that achieved in the current study. A current large-scale study by the ManyBabies consortium which aims to test the predictions made by Hunter and Ames [26] may give insight into this possibility [31]. Such subtle effects would not be consistent with the claims of larger familiarity preferences that have appeared in the literature, however.

### 5.4 Formal theories of learning and attention

The lack of familiarity preferences in the current study can also be attributed to the particular learning problems participants are solving: whether familiarity preferences occur can depend on the particular learning context and the particular learning problem. This context-dependence is reflected in meta-analyses investigating familiarity preferences across paradigms. For example, when tested on word segmentation in their native language, infants showed preferences for familiar stimuli throughout the first year [5]. In contrast, when tested on statistical learning of novel words, infants showed consistent preferences for novel stimuli, from 4 months to 11 months of age [6].

The seemingly contradictory results on the direction of preferences in infant research highlight the need for formal theories of how the learning problem influences attention. Dubey and Griffiths [15] gave an example of such a formal account by showing that when past and present events are correlated, rational agents, under some assumptions, develop a tendency to attend to familiar stimuli to prepare for the most likely future events, while in uncorrelated environments, novelty preferences are optimal. Ideal learners attempting to maximize their expected information gain consistently seek novelty when trying to learn a single concept. Formal models of the learning problem being solved in experiments may therefore give more principled and precise predictions beyond identifying factors like “prior exposure” as determinants of attentional preferences.

Importantly, RANCH’s predictions outlined in this paper apply to the specific case when the learner makes noisy observations of stimuli in a perceptual space, updates their beliefs about the location of a single concept in that space, and makes decisions about whether to continue observing based on their expected gain of information relative to this belief updating process. The modular structure of RANCH allows for each of these components (perceptual embedding space, learning model, and linking hypothesis) to be modified; such modifications could lead to different predictions. For example, the embedding space could be chosen to more closely resemble a psychologically validated perceptual space [e.g. 33], though in this case, we found that such alternatives did not affect embedding structure qualitatively (Supplemental Figure 12). Similarly, the simple concept learning model we used could be modified to account for more complex, or more general, representation learning [28, 50]. For example, we could model our participants as learning multiple concepts, deciding whether each stimulus is part of the same concept or a new concept.

Another promising direction is to explore RANCH’s applicability to finer timescales of looking behavior, enabling a more detailed examination of within-trial fluctuations in attention. Recent work suggests that analyzing moment-by-moment dynamics can help disentangle distinct learning mechanisms [43].Since RANCH models decision-making at the level of individual perceptual samples, it is well-suited to capture these fine-grained attentional shifts.

### 5.5 Conclusion

From birth, infants actively explore their environment through selective attention. Looking time, a widely used measure in developmental psychology, has long been used to make inferences about infants’ conceptual and perceptual capacity. Here, we measured nuances of key looking time phenomena, such as habituation and dishabituation, by varying familiarity with a stimulus prior to dishabituation (Experiment 1), as well as the similarity between the familiar and novel stimulus (Experiment 2). We found that these behavioral nuances are well captured by RANCH, an image-computable, rational model that learns from noisy perceptual samples and decides how many samples to take based on their expected information gain. Beyond modeling infant looking during perceptual learning in particular, our work provides a general framework that instantiates and tests hypotheses about the computations underlying infant looking.

## 6 Methods and Materials

### 6.1 Model specifications

#### 6.1.1 Perceptual representations

We derived principled low dimensional representations of our stimuli from ResNet-50 [24]. We ran a Principal Component Analysis and used the first three principal components. This decision was made due to the computational demands of our model. Each added dimension would increase the total run time exponentially. The first three components captured 57.9% of the variance.

Given that we were using discrete grid approximation in our inference, the model was sensitive to the absolute values of the embeddings. When these embeddings values were close to or exceeded the range of the approximation grids, this would cause bias in our simulation result. We therefore scaled our embeddings down to fall squarely into our grid. Simulations have shown that embedding that were too large would result in biased model behaviors (Supplementary Information, Figure 10). Since embedding distances were larger for stimulus type experiment, especially for animacy violations, the downscaling had to be more extreme.

#### 6.1.2 Linking hypotheses

We compared two additional information theoretic measures with Expected Information Gain (EIG): Kullback– Leibler divergence and Surprisal. Each of these measures has been invoked in the literature as a plausible account for explaining looking behaviors across developmental stages [KL divergence: 41, surprisal: 30].

Our key linking hypothesis, EIG, is a forward-looking measure. It is computed as the product of the posterior predictive probability of the next perceptual sample (*p*(*z*_*t*+1_ |*θ*_*t*_)) and the information about the concept *θ* gained conditioned on that next hypothetical sample, using a grid of possible subsequent samples(s): 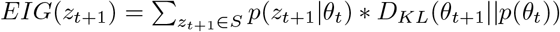

Both KL-divergence and surprisal are backward-looking measures. The KL divergence at any given point is calculated by the posterior distribution after seeing the current perceptual sample and before seeing the current perceptual sample: 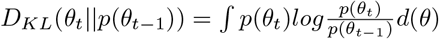

Surprisal is calculated as the negative log likelihood of the current perceptual sample under the prior distribution: *surprisal*(*z*_*t*_) = *−log*(*p*(*z*|*θ*_*t−*1_))

We ran individual simulation and parameter search for each of the three linking hypotheses to ensure fair comparison.

#### 6.1.3 Scaling procedure for developmental interpretation

To find the best fitting parameters for each age group we employed a strategy we called “joint scaling”. To evaluate model fit, we first mapped model samples onto looking time by finding the best linear scaling of model samples in a regression of the form `LT ∼ model_samples.` The parameter set that minimized the RMSE between the scaled model samples and looking time was what we considered the best parameter set. In the case of “joint scaling”, we regressed both the adult and the infant simulations against their looking time simultaneously, to find how varying the infant and adult parameters affected the fit of the scaled model samples to the adult and infant data. This joint scaling procedure put pressure on the parameters, rather than the scaling, to account for the fact that adults’ looking time were a lot shorter than infants’ looking time.

The interpretable parameters of RANCH were the priors on *µ* and *σ* and the priors on the noise. The priors on *µ* and *σ* were parameterized by a normal inverse-gamma prior with *µ*_*prior*_, *ν*_*prior*_, *α*_*prior*_ and *β*_*prior*_, the conjugate prior to a normal distribution with unknown mean and variance. While the mean was fixed to be 0 (*µ*_*prior*_), variation in the other parameters expressed distinct hypotheses about the precision of the location of the concept (*ν*_*prior*_), as well as the variance of the concept (*α*_*prior*_ and *β*_*prior*_). The learner’s prior on noise was parameterized by zero-mean Gaussian distribution with standard deviation (*σ*_*epsilon*_).

### 6.2 Experimental procedure

Below we described in detail the experimental procedure. We used Experiment 1 to denote the experiment that manipulates exposure duration, and Experiment 2 to denote the experiment that manipulates stimuli similarity.

#### 6.2.1 Infant experiment

##### 6.2.1.1 Participants

For Experiment 1, we tested a combined sample of 103 7-10 month old infants, with 31 in the first sample, 35 in the second sample and 37 in the third sample (*M*_*age*_ = 9.38 months, 47 female). 10 participants were excluded completely due to fussiness. An additional 58 individual trials were excluded.

For Experiment 2, we tested a combined sample of 123 7-10 month old infants, with 57 collected via Zoom (7 complete exclusions, 42 trial exclusions), 66 collected via CHS (3 complete exclusions, 26 trial exclusions).

Complete or trial exclusions occurred because (1) infants fussed out of the experiment at an earlier stage of the experiment, (2) infants looked at the stimuli for less than a total of 2 seconds, (3) there were momentary external distractions in the home of the infant or (4) the gaze classifier performed so poorly that manual corrections were not viable.

##### 6.2.1.2 Stimuli

In Experiment 1, we presented infants with a series of animated animals, created using “Quirky Animals” assets from Unity (Figure 3; link to assets). The animals were walking, crawling or swimming, depending on the species. In a previous study not included here (but see OSF repository), we showed infants a different stimulus set and failed to elicit replicable habituation, novelty or familiarity preferences. As a result, we modified the stimuli to be more appealing. We also separated exposure trials with curtains opening and closing to accentuate exposure trials as separated occurrences of the stimulus.

In Experiment 2 (Figure 2A), we introduced animacy violations using the “3D Prop Vegetables and Fruits”(link to assets) assets from Unity and animated bouncing and rotating trajectories for them. To create number violations, we duplicated the videos of single animal Unity assets and pasted them side-by-side.

##### 6.2.1.3 Procedure

Data collection was performed primarily synchronously on Zoom, except for the CHS replication of Experiment 2. In all cases, infants were recruited from CHS [47] and Facebook. Parents were instructed to find a quiet room with minimal distractions, place their child in a high chair (preferred) or on their lap, and to remain neutral throughout the experiment. Infants were placed as close as possible to the screen without allowing them to interact with the keyboard.

Both main experiments followed a block structure, where each block was divided into two sections: (1) a exposure phase and (2) a test event. Each block was preceded by an “attention getter”, a salient rotating star. During the exposure period, the infant was familiarized to a particular animal in a series of exposure trials. Each exposure trial was a 5 second sequence: curtains opened for 1 second, the animated animal moved in place for 3 seconds, and then the curtains closed for 1 second. In Experiment 1, the number of exposure trials (the “exposure duration”) varied between blocks. In Experiment 2, the number of exposure trials was always 8 or 9.

During the test event, the infant saw either the same familiar animal again, or a novel stimulus. The onset of the test event was not marked by any visual markers, but a bell sound played as the curtains opened, to maximize the chance of engagement during the test event. On Zoom, the test event used an infant-controlled procedure: the experimenter terminated the trial when the infant looked away for more than three consecutive seconds. On CHS, test trials would always be played for 40 seconds. In both cases, looking time was then defined as the total time that the infant spent looking at the screen from the onset of the stimulus until the first two consecutive seconds of the infant looking away from the screen. If the infant did not look away after 60 seconds of being presented with the test event, the next block automatically began and infants’ looking time for that test event was recorded as 60 seconds.

In Experiment 1, the novel stimulus was always another animal. In Experiment 2, novel stimuli violated one of four properties of the familiar animal: pose, identity, number or animacy.

In Experiment 1, each infant saw six blocks: Three different exposure durations ({0,4,8}, {1,3,9} or {2,4,6}) appeared twice each, once for each test event type (familiar or novel). The longer exposure durations (8 or 9) were chosen based on our previous pilot studies with a different stimulus set (OSF repository), and the shorter durations were chosen to provide limited learning experience with the familiar stimuli. The order of blocks was counterbalanced between infants, and pairs of animals (familiar and novel) were counterbalanced to be associated with each block type. Which animal was shown as the familiar and which as the novel (if it was a novel test trial) was randomized in each block.

In Experiment 2, each infant saw four blocks within a session - the familiar trial was always one of the four test trials, and the remaining three test trials were picked randomly from the remaining four violation types (pose, identity, number, animacy).

The control experiment showed six standalone trials: Infants saw two single animals, two pairs of animals and two single vegetables, to measure baseline interest in these stimulus categories.

##### 6.2.1.4 Looking time coding

To code the infants’ looking time we used iCatcher+, a validated tool developed for robust and automatic annotation of infants’ gaze direction from video [16]. To obtain trial-wise looking time, we merged iCatcher+ annotations with trial timing information, thereby fully replacing manual coding of looking time. The correctness of iCatcher+ coding was manually supervised to check for failed face detections or distractions.

##### 6.2.1.5 Statistical analysis

We ran the pre-registered statistical analyses for both experiments. All statistical analyses were ran in R using the package `lme4` [4].

For Experiment 1, we ran a lme4 model with the following specification: `log(looking time) ∼ poly(exposure_duration, 2) * test_type [identity or familiar] + log(block_number) + (1|subject).` The `exposure duration` term referred to the amount of prior exposure received by the infant (0 to 9 prior exposures), and the second polynomial tested for non-linearity in the effect on looking time, as predicted by some theories of infant attention. This non-linearity could occur primarily for looking to the familiar stimuli (Figure 1A), or both familiar and novel stimuli (Figure 1B), hence the interaction with `test type.` We controlled for block number to account for general decreasing interest in our stimuli as the experiment goes on.

For Experiment 2, we ran a linear mixed effects regression, again predicting looking time, using violation type as the main predictor: `log(looking_time) ∼ test_type [animacy, pose, identity, number, familiar] + log(block_number) + (1|subject).` Here, we set familiar as the reference level for `test type`, and asked whether the other test types were significantly different from that familiar reference level. We again controlled for block number.

In our pre-registered model comparison analysis, we used a chi-square test to compare the above regression model above to a regression model in which test type only had two levels, familiar or novel.

#### 6.2.2 Adult experiment

##### 6.2.2.1 Participants

Both experiments were run on Prolific (Experiment 1: N = 522; Experiment 2: N = 536). Two experiments had the same pre-registered exclusion criteria. We excluded participants if (1) the standard deviation of their reaction time across all trials was less than 0.15 (indicating key-smashing, Experiment 1: N = 0; Experiment 2: N = 0), (2) spent more than three absolute deviations above the median of the task completion time as reported by Prolific (Experiment 1: N = 47; Experiment 2: N = 43), and (3) provided the wrong response to more than 20% of the memory task (Experiment 1: N = 13; Experiment 2: N = 33). A total of 52 and 68 participants met at least one of the three criteria and were excluded from the final analysis for Experiment 1 and Experiment 2, respectively. We also excluded a trial if the trial was three absolute deviations away from the median in the log-transformed space across reaction time from the remaining participants.

The final sample included 470 participants for Experiment 1 and 468 participants for Experiment 2 (Experiment 1: *M*_*age*_ = 33.0 years, *SD* = 12.3; Experiment 2: *M*_*age*_ = 31.8 years, *SD* = 11.3).

##### 6.2.2.2 Stimuli

The stimuli used in adult experiment were identical to the ones used in infant experiment. In Experiment 1, the stimuli were selected from the animal unity set. In Experiment 2, both the animal and vegetable sets were used.

##### 6.2.2.3 Procedure

Procedurally, Experiment 1 and Experiment 2 were similar. Both were self-paced visual presentation studies. Participants were instructed that they would be looking at a sequence of animated stimuli at their own pace. At each trial, they can press a key on the keyboard to move on to the next trial after a minimum viewing time of 500 ms. Each study consisted of multiple blocks. Each block consisted of multiple trials. Between blocks, participants answered a simple memory question (“Have you seen this animation before?”). This memory question was used as an attention check.

Each block contained multiple trials of one stimulus being repeatedly presented (the familiar trials) and one trial that showed a different stimulus (the novel trial). The novel trial always appeared as the last trial. In Experiment 1, the total number of trials ranged from two to eleven. The novel trial was randomly drawn from the stimuli set that researchers haven’t seen before. In Experiment 2, the block consisted of either two, four, or six trials. There were a total of eight types of stimuli in the familiar trials. They were all combinations of the three features that each include two levels: animacy (e.g. animate or inanimate), number (singleton or pair), and pose (facing left or facing right). The novel trial can differ from the familiar trials in one of the four dimensions: pose, number, identity, and animacy. Both experiments also had blocks that only include one stimulus (the familiar block).

The order of the block was semi-randomized in both experiments. Experiment 1 has 15 blocks in total. We grouped the total number of trials in each block into five pairs of two: {2, 3}, {4, 5}, {6, 7}, {8, 9}, {10, 11}, and randomly sampled 1 block length from each pair. In other words, each participant would receive 5 different lengths of repeating familiar trials, stratified by 5 levels. We controlled the distribution of the block lengths by making sure the first half and the second half of the experiment each had the same number of blocks with different block lengths. Experiment 2 had 24 blocks in total. We grouped the twenty-four blocks into four groups. Each group consisted of two familiar blocks and one block from each of the four novelty types. The order of blocks within each group was randomized.

##### 6.2.2.4 Statistical Analysis

We ran the pre-registered statistical analyses for both experiments. All statistical analyses were ran in R using the package `lme4` [4].

For experiment 1, we ran a linear mixed effect regression predicting the log transformed of looking time with the following specification: `log(looking time) ∼ trial_number + is_first_trial + trial_number * trial_type + log(block_number) + (trial_number * trial_type | subject) + (is_first_trial + trial_number | subject)`. This model allowed us to investigate whether the number of familiar trials has an impact on the looking time (i.e. habituation), and whether the trial type (familiar or novel) mediated the influences. This model failed to converge. Following the pre-registered procedure, we pruned the model to include only by-subject random intercept. The pruned model suggests that there is evidence for habituation and dishabituation.

For experiment 2, we ran a similar model and incorporated the effect of different novelty type. The particular model specification is as follows: log(total_rt) ∼trial_number + is_first_trial + (trial_number + is_first_trial) * stimulus_number + (trial_number + is_first_trial) * stimulus_pose +(trial_number + is_first_trial) * stimulus_animacy + (trial_number + is_first_trial) * novelty_type + log(block_number). novelty_type had five levels, including the familiar trial and four types of violation. This model can help us test two hypotheses: (1) whether our experimental paradigm captured habituation and dishabituation and (2) whether the magnitude of dishabituation was influenced by the similarity between the novel trials and the familiar trials.

## 7 Supplementary Information

### 7.1 Deriving predictions for Goldilocks model

Below is an illustration of how predictions for a habituation/dishabituation experiment were derived for the Goldilocks model. In all cases, as infants are familiarized, the surprisal of the familiar stimulus goes down (Blue arrows pointing left), and the surprisal of the novel stimulus has to go up so that probabilities assigned to events add up to 1 (Red arrows pointing right). Depending on the assumed surprisal of the exposure stimulus, this model predicts three qualitatively different trajectories of preferences to familiar vs. novel stimuli as a function of prior exposure: (1) If the assumed surprisal is below the optimal level to maximize attention, the model predicts an increase, and then a decrease in novelty preferences. (2) If the assumed surprisal is at the optimal level, then no preference is predicted. (3) If the assumed surprisal is higher than optimal surprisal, the model predicts an increasing, and then decreasing familiarity preference.

In our work, we assumed a scenario consistent with the first assumption – i.e. that the initial surprisal of stimuli is below the optimal level. This is the only regime in which novelty preferences arise, and therefore had the highest chance of agreeing with with our data, as well as predictions made by Hunter and Ames [26].

### 7.2 Parameter robustness

To evaluate RANCH’s robustness to different parameters, we tested its fit for all possible parameter settings. Performance across the 162 parameter settings was relatively stable, yielding a moderate range RMSE values for infants and adults across linking hypotheses. In both cases most parameter settings outperformed the fit achieved by baseline models (Table **??**). The distribution of RMSE values across parameter settings are shown in the histograms below (Figure 9).

**Figure 9:**
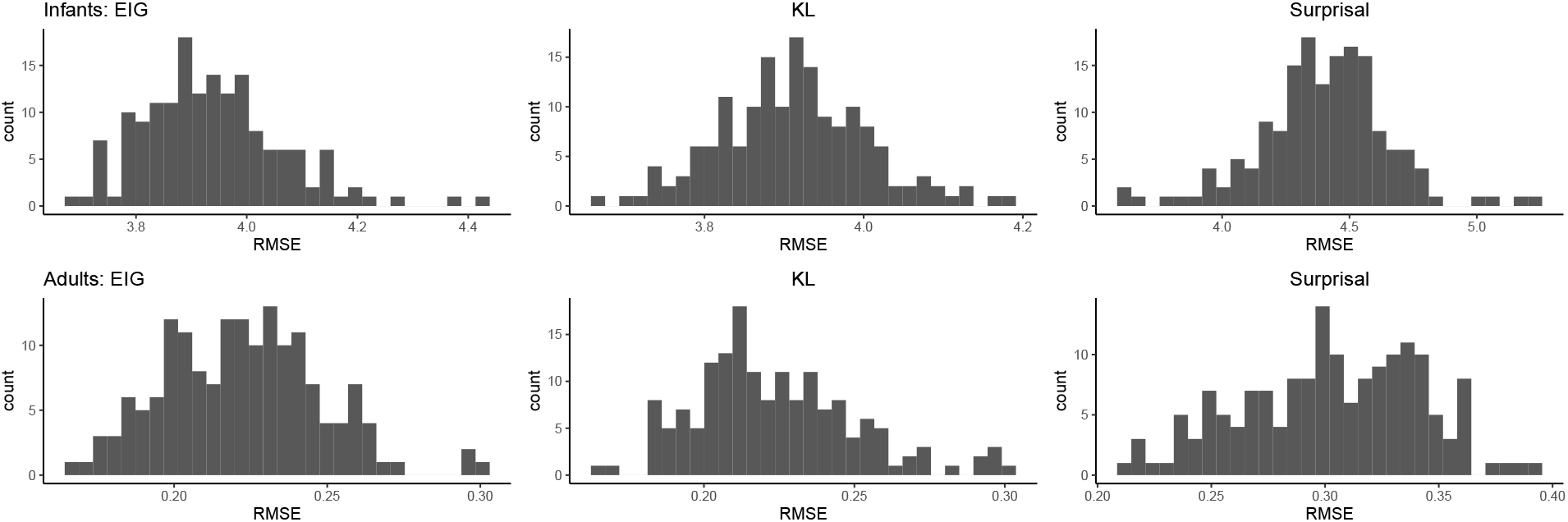
Histograms of RMSE for different linking hypotheses show that RANCH is largely robust to variations in parameters.

**Figure 10:**
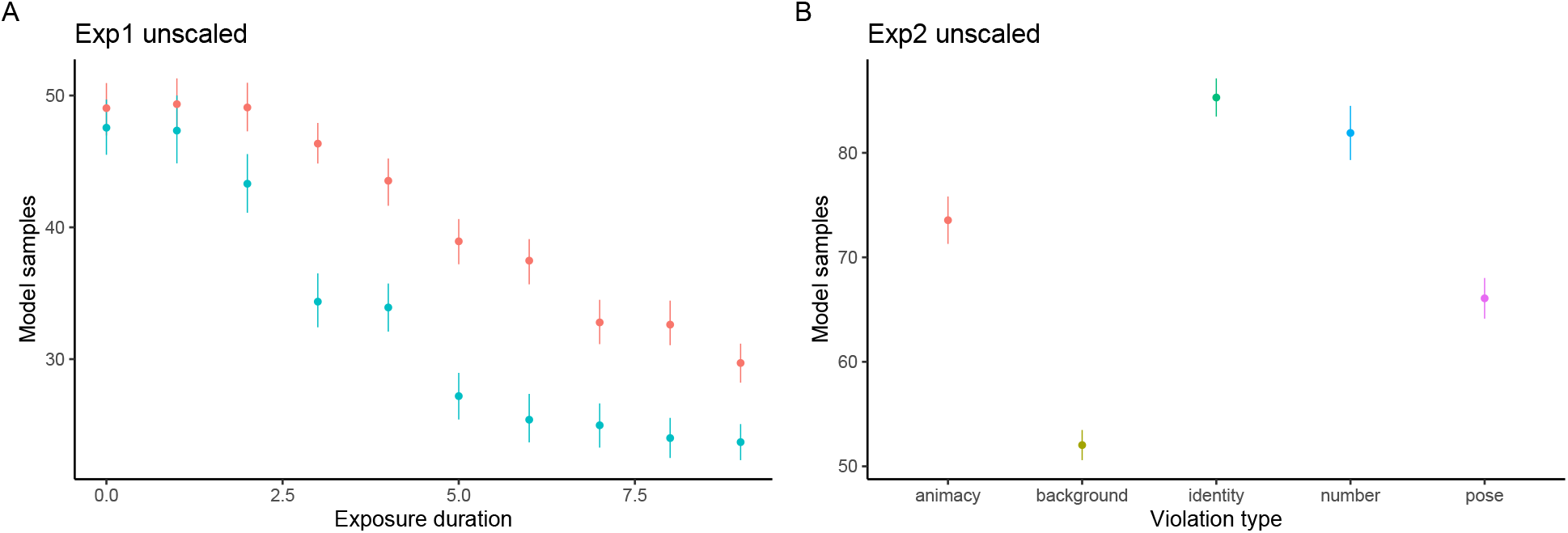
The effects of scaling on model results. (A) Experiment 1 with unscaled embeddings resulted in a reversal of familiar vs. novel. (B) Experiment 2 with unscaled embeddings resulted in an unintuitive ordering.

**Figure 11:**
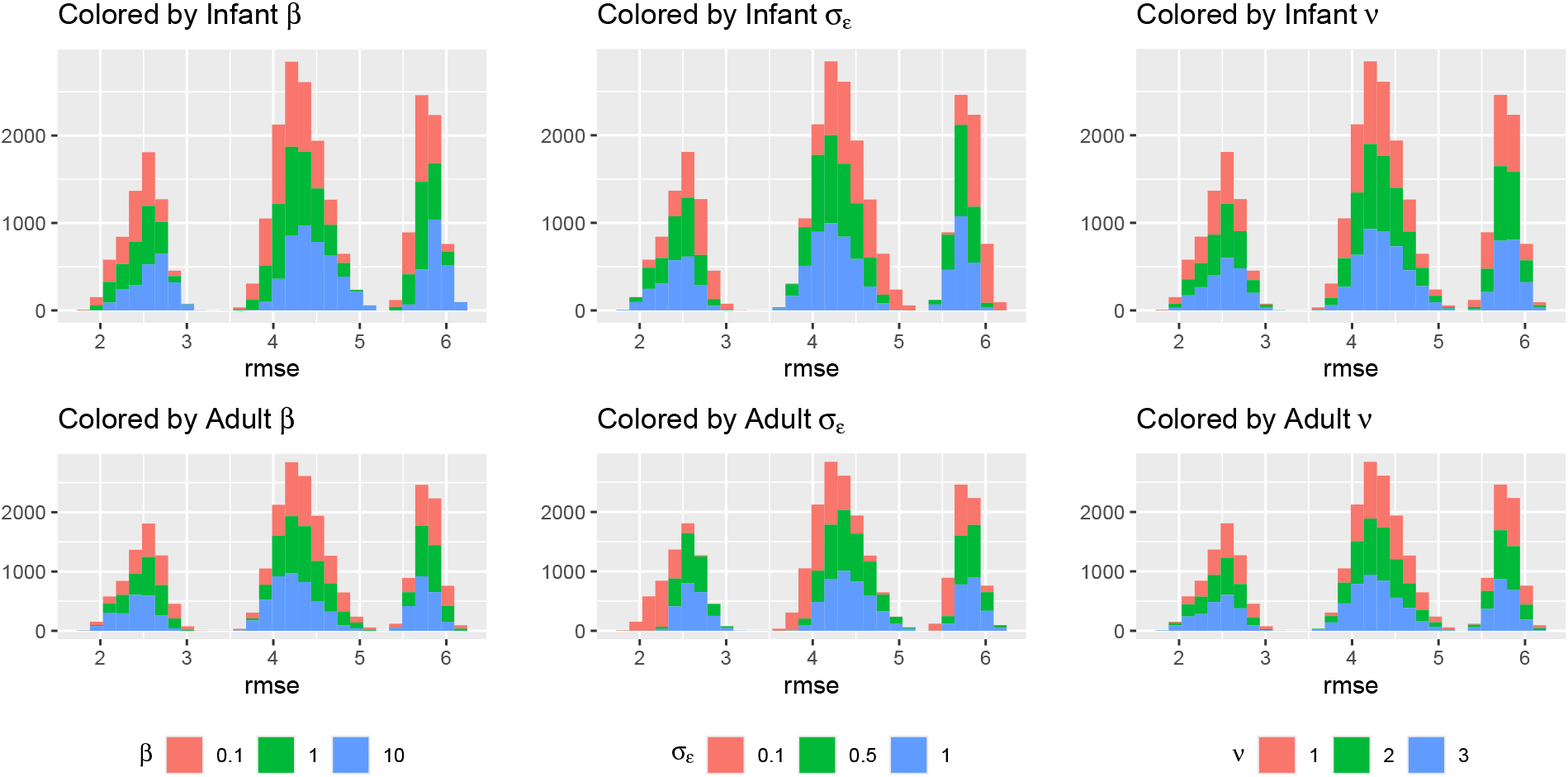
Histograms of model fit colored by priors. X-aixs shows the RMSE of the model fit with the behavioral data. Results suggest that when the learner’s noise is high, RANCH fits infant data better.

**Figure 12:**
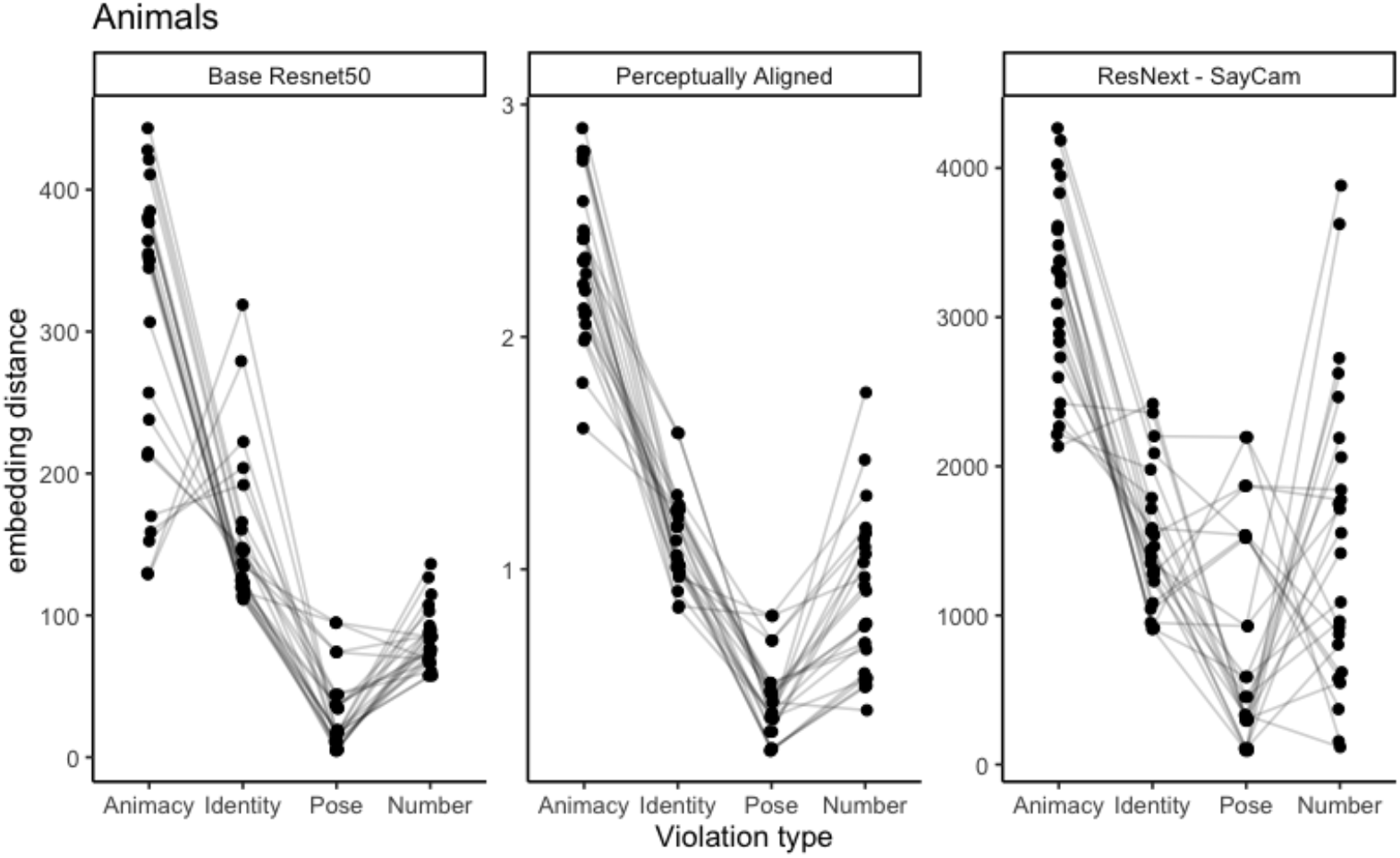
Embedding distances of different violation types for (1) a base ResNet50 model, (2) a perceptually aligned embedding model [33]. and (3) a ResNet50 model trained on SAYCAM data [39]

**Figure 13:**
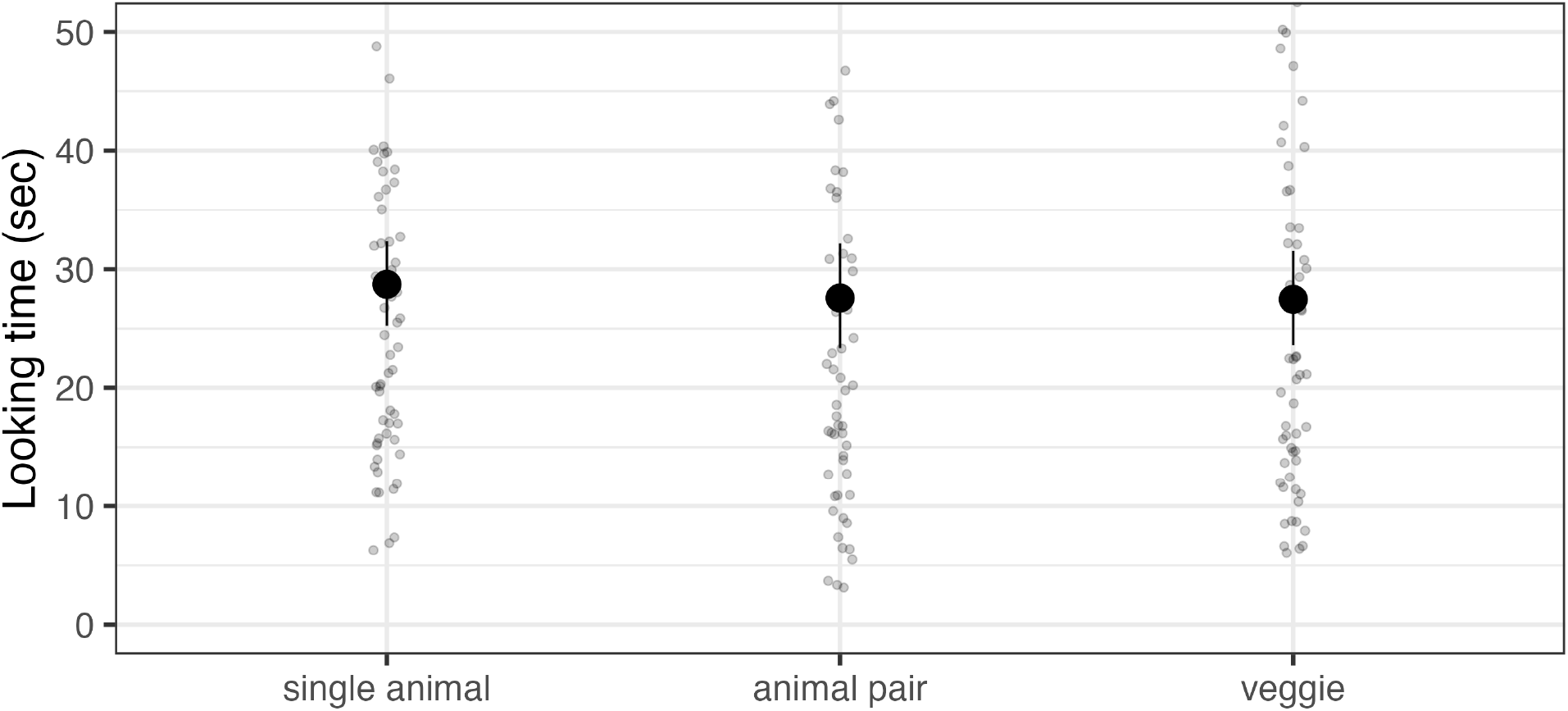
Control experiment for Experiment 2 (infants), checking for baseline differences in interest. We found no significant differences in baseline looking.

### 7.3 Scaling the embedding space

In order to achieve sensible results, it was important to bring the stimulus embeddings in line with the approximation grid used for inference. Below we show what happens when embeddings exceeded or came close to the edges of the our approximation grids. In Experiment 1, unscaled embeddings resulted in a reversal of familiar vs. novel model samples, such that the model takes more samples from the familiar than the novel. In Experiment 2, unscaled embeddings resulted in an unintuitive ordering which is not in line with the embedding distances (Figure 10).

### 7.4 Developmental comparison of parameter fits

A naive approach to comparing parameter fits between infants and adults would be the following: After joint scaling, pick the set of parameters for infants and adults that together minimized RMSE. If we do this, we obtain the following parameter sets:

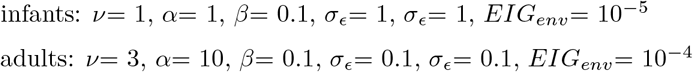

If we compare these two sets of parameters to each other, and interpret these differences, we may come to the following conclusion: RANCH fits best when infants’ *σ*_*ϵ*_ is higher than that of adults, and *β*_*prior*_ is higher than that of adults, suggesting that infants both expect their perception to have more noise, and that infants have more prior uncertainty about the size of the standard deviation.

However, a direct comparison between these parameter sets does not give insight into the relative stability of RANCH’s fit to the data if we change any of these parameters: It may be that if a parameter was slightly different, it would not make a big difference for model fit.

Thus we checked the sensitivity of the model fit to changes in the parameters. To do so, we plotted RMSE histograms colored by the value of a certain parameter, from a certain age group. Here we see that the overall modes of the RMSE histograms are dominated by world EIG, likely because this parameter is the main threshold that can account for the difference in scale between infant and adult looking time. However, of the other parameters, we see that RMSE only shows sensitivity to *σ*_*ϵ*_. Other parameters that are different between the best fits are not stable across settings. We therefore conclude that the main interpretable parameter difference is in *σ*_*ϵ*_, the learner’s noise prior, being higher in infants than in adults.

### 7.5 Embedding distances of different violations categories, by embedding model

We chose ResNet-50 as our embedding space, but there are other embedding space available. Below we plotted the embedding distances of the different violations. The lines connect a single stimulus, and the y-axis represents the embedding distances to all the stimuli that would constitute a certain violation. For example, for a single animal stimulus, the y-value for ‘animacy’ corresponds to the distance between the single animal and all the single fruit/vegetable stimuli. Since the embedding distance between different categories were similar to each other, we did not run the full simulations on each of the embedding space.

### 7.6 Control experiment for Experiment 2 in infants

Given that the exposure phase always consisted of a single animal, it is possible that differences in looking time during test trials of different types were a result of differences in interest in the stimuli themselves (e.g. our inanimate stimuli may have been intrinsically more interesting than our animals). To test whether infants were differently interested in our stimuli due to intrinsic stimulus properties, we ran a control experiment in which we showed infants our novel stimulus types without any prior exposure. Infants saw three types of stimuli for as long as they wanted, up to 60 seconds: single animals, fruits/vegetables, or pairs of animals. There were six trials per session, so infants saw two of each stimulus type. We tested 35 infants for this experiment, with 2 complete subject exclusions and 10 single trial exclusions.

We found no evidence for differences in baseline interest in our stimuli: neither looking to inanimate stimuli nor animal pairs differed significantly from looking to single animals (inanimate stimuli: *β* = -0.02; *SE* = 0.08; *t* = -0.31; *p* = 0.758; animal pairs: *β* = -0.11; *SE* = 0.08; *t* = -1.38; *p* = 0.169). In sum, infants showed equal interest in all categories of test stimuli, prior to habituation; the condition differences observed in Experiment 2 must derive from the cognitive processes of habituation and dishabituation.

## Footnotes

1 ResNet-50 is by no means the only possible method for deriving perceptual representations; instead it is a simple, standard choice. We also explored alternative models, such as a CNN trained on egocentric videos from the child’s perspective and CNN whose representations were aligned to human categorization behavior [38, 33]. However, we did not find substantial differences between the embedding spaces provided by ResNet-50 and other (often more complicated) models (Supplementary Information, Figure 12).

2 Note that we pre-registered a significant effect of identity violation on looking time as a data quality check, since we had replicated that effect in both infant experiments. However, surprisingly, the identity condition was the only condition that differed from the Zoom experiment, so here we deviated from pre-registration and report findings from this sample given the high value of having a second sample of infant data.

